# Structure of HK97 small terminase:DNA complex unveils a novel DNA binding mechanism by a circular protein

**DOI:** 10.1101/2023.07.17.549218

**Authors:** Maria Chechik, Sandra J. Greive, Alfred A. Antson, Huw T. Jenkins

**Affiliations:** York Structural Biology Laboratory, Department of Chemistry, University of York, York YO10 5DD, United Kingdom

**Keywords:** HK97 small terminase, bacteriophage, protein:DNA complex, cryoEM structure

## Abstract

DNA recognition is critical for assembly of double-stranded DNA viruses, in particular for the initiation of packaging the viral genome into the capsid. DNA packaging has been extensively studied for three archetypal bacteriophage systems: *cos*, *pac* and phi29. We identified the minimal site within the *cos* region of bacteriophage HK97 specifically recognised by the small terminase and determined a cryoEM structure for the small terminase:DNA complex. This nonameric circular protein utilizes a previously unknown mechanism of DNA binding. While DNA threads through the central tunnel, unexpectedly, DNA-recognition is generated at its exit by a substructure formed by the N- and C-terminal segments of two adjacent protomers of the terminase which are unstructured in the absence of DNA. Such interaction ensures continuous engagement of the small terminase with DNA, allowing sliding along DNA while simultaneously checking the DNA sequence. This mechanism allows locating and instigating packaging initiation and termination precisely at the *cos* site.

## Introduction

The large group of dsDNA viruses comprising tailed bacteriophages and herpesviruses assemble by packaging their DNA into preformed procapsids. The key component ensuring specific recognition of bacteriophage DNA from the mixture of all the nucleic acids contained in the host cell, and initiating its packaging is the small terminase protein. The nascent genome is usually produced as multiple copies, joined head-to-tail, in long concatemers of dsDNA. Packaging initiation occurs when the small terminase binds to a specific sequence (*pac* or *cos*) in the genomic DNA, facilitating the cleavage of the dsDNA by large terminase and subsequent assembly of the large terminase bound DNA complex onto the portal protein vertex of the capsid for packaging to begin^1, 2, 3^.

Termination of packaging must only occur after a complete copy of the genome has been inserted into the capsid and subsequent cleavage of the DNA by the nuclease domain of large terminase is achieved by one of two methods. In the case of headful (*pac*) packaging, termination cleavage occurs randomly after approximately 102-105% of viral genome has been packaged into the virion. In contrast for *cos* viruses, such as λ or HK97, termination cleavage occurs precisely at the boundary of the next genomic copy, defined by the *cos* sequence, and requires small terminase^4, 5^. After termination cleavage, the large terminase motor:DNA complex is transferred to a new empty capsid where it continues packaging the next copy of the genome in the concatemeric DNA. This transfer of the packaging complex happens between 2 to 12 times per concatemer, depending on the phage^6, 7, 8^.

Although structural information is already available for small terminases from more than 10 dsDNA bacteriophages (**Supplementary Fig.** 1^4, 9, 10, 11, 12, 13, 14, 15, 16, 17, 18, 19, 2^, unpublished pdb:2ao9,4xvn,6ejq) the mechanism of DNA recognition and binding remains the subject of a major debate. All small terminase structures broadly resemble each other in architecture, with 8-12 protomers oligomerized in a ring to form a central channel in the middle, with the N-terminal segments arrayed around the outside. The C-terminal region is often disordered^4, 12, 18, 19, 20^. While many of the N-terminal segments are folded into helix-turn-helix (HTH) -like folds presumed to be the DNA-binding domain (DBD), and sufficient for DNA binding in λ phage and SF6^10, 21^, the observed structural variation has confounded development of a universal consensus for the mechanism of small terminase binding to DNA. However, based on the available data, two alternative models have been proposed: a wrapping model where the DNA wraps around the outside of the circular oligomer by interacting with the N-terminal DBDs and a threading model where the DNA is threaded through the internal channel of the circular oligomer^3^. However, despite being the focus of research for many years, no structure of a DNA-bound small terminase complex has been determined.

*Escherichia coli* bacteriophage HK97 has been the subject of intensive research, and along with λ and P22^17, 21, 22^, serves as an excellent model system for understanding the mechanism of virus assembly and DNA packaging. Recent work on HK97 developed an *in vitro* packaging system and determined the structures for both large and small terminases^4, 23^. In this work we defined the boundaries of the minimal sequence within the *cos* region that is specifically recognised by the small terminase. We also report the structure of the terminase:DNA complex, determined by cryoEM, providing the first experimental structural information for the small terminase:DNA complex.

## Results

### Specific small terminase binding site within the HK97 *cos* region

Preliminary data indicated that the recognition site for small terminase lay between positions -80 and +472 spanning the cleavage site (between -1 and +1) in the *cos* region^24^. To define the binding site more precisely, we used progressively shorter DNA oligos encompassing the *cos* cleavage site to probe for small terminase:DNA binding by electrophoresis mobility shift assays (EMSA) using fluorescently labelled DNA (**Supplementary Fig. 2**). Subsequently, unlabelled DNA oligos were used to locate the boundaries of the binding site with single base pair resolution (**Fig. 1**). The minimal region necessary for small terminase binding comprises the 15bp (position 15-29) starting from 15 bp downstream of the *cos* cleavage site.

**Figure 1.**
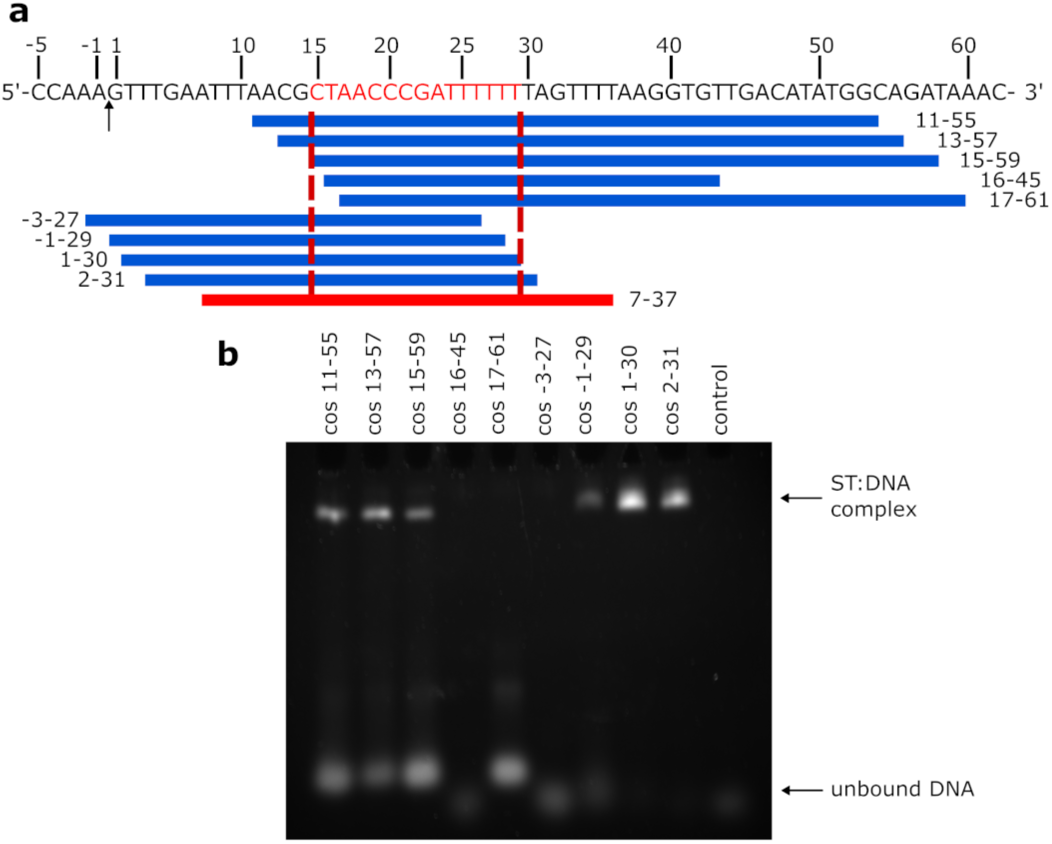
DNA binding site of small terminase. **a** Sequence and schematic diagram of HK97 *cos* site: arrow indicates cleavage site (between position -1 and 1) during genome packaging. Red segment (15-29) - essential DNA binding site. Blue blocks represent different dsDNA fragments. Red block (7-37) - oligo that was used for complex formation. Red dotted lines show the border of DNA binding site. **b** Electrophoretic mobility shift assay (EMSA) of small terminase with dsDNA. Control lane – no protein.

### CryoEM structure of the small terminase:DNA complex

After determining the minimal small terminase binding site, EMSA was used to confirm that dsDNA fragments of several different lengths: 21, 25 and 31 bp, all containing this binding site in the centre of the fragment, were sufficient to enable formation of a complex with small terminase (**Supplementary Fig. 3**). Of these fragments, the longest, with 31 bp, was selected to form a small terminase:DNA complex suitable for structural determination by single particle cryoEM. The complex was purified from unbound DNA using size exclusion chromatography (**Supplementary Fig. 4).** The structure of the small terminase:DNA complex was determined at a resolution of 3.0 Å by cryoEM (**Fig. 2, Supplementary Fig. 5-9, Supplementary Tables 1&2**). Unexpectedly the complex is asymmetric with the DNA, threaded through the central channel, bent towards one side of the small terminase nonamer.

**Figure 2.**
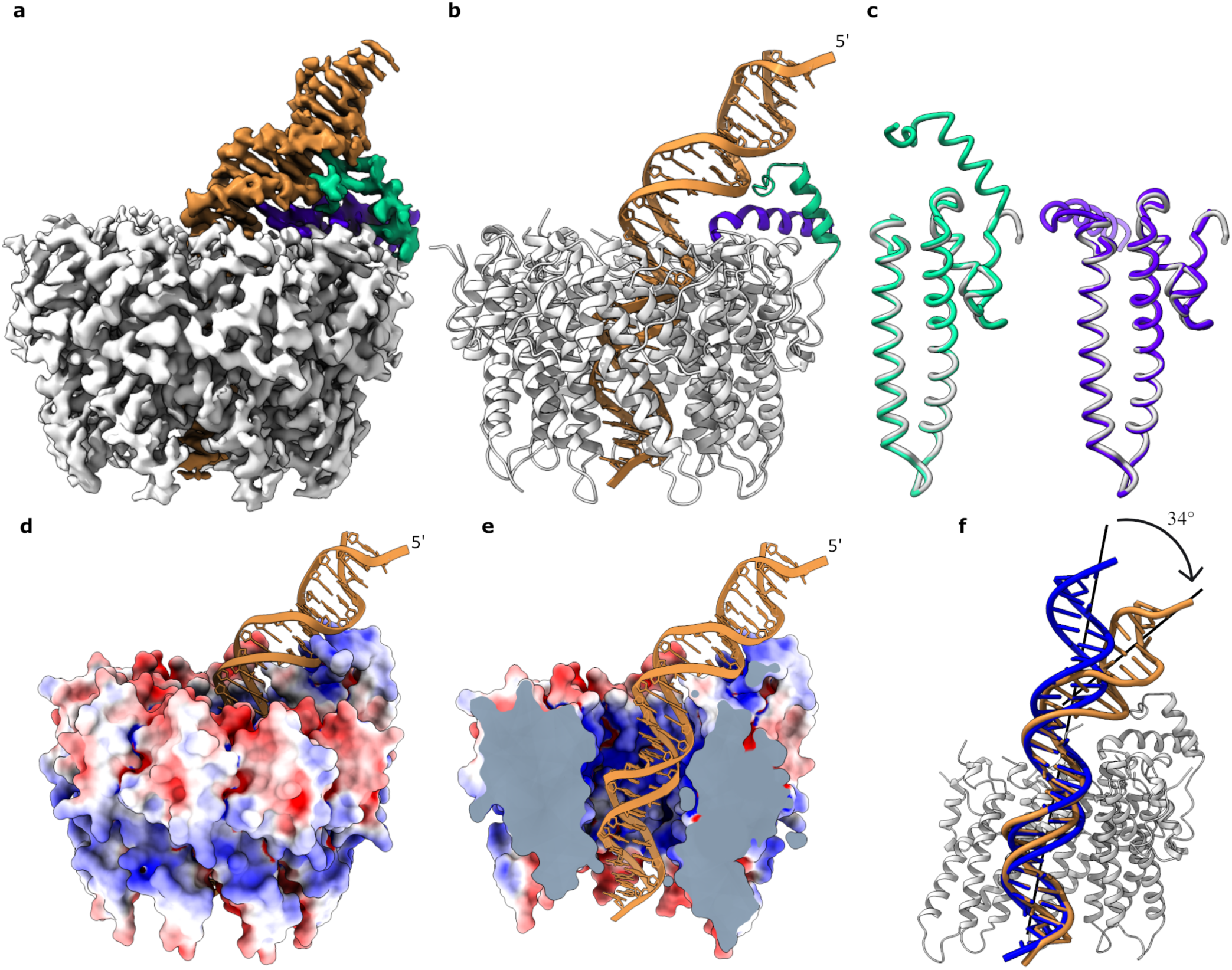
Structure of small terminase:DNA complex. **a** CryoEM map and **b** ribbon representation of small terminase:DNA complex. Nine protomers (grey) form the nonameric structure of small terminase. N-terminal helix from chain A (green) and a C-terminal helix from chain B (purple) form a DNA binding motif that directly interacts with DNA (gold). **c** Superposition of one monomer (grey) from the previously determined X-ray structure of small terminase (PDB 6z6e) with chain A (green) and B (purple) from the cryo-EM structure. **d** Molecular surface of the small terminase coloured according to electrostatic charge, with DNA shown as ribbon. Negative and positive charges are in red and blue respectively, varying from −5 to 5 kT/e. **e** Cross section of the map showing the electrostatic surface charges (calculated as in **d**) inside the channel. **f** Alignment of DNA (gold) with ideal B-form DNA of the same sequence (blue). Mainchain alignment of 11 base pairs from the bottom end of the DNA was performed with LSQ in Coot.

As observed in the previously reported crystal structure^4^ the small terminase oligomer contains 9 protomers surrounding a central channel. In the crystal structure the diameter of the channel at its narrowest point is 18 Å whilst in the cryoEM structure the channel is ∼3% larger. While part of this difference in size may reflect the accuracy of the calibrated pixel size for the cryoEM structure, it is possible that the channel expands slightly to accommodate DNA. In the cryoEM structure residues Asp24 - Asp124 are well defined for all nine protomers (**Fig. 2a,b**). The mainchain of this region in the cryoEM structure aligns to the crystal structure with a r.m.s.d. of 0.5 Å (**Fig. 2c)** indicating that binding to DNA does not change the conformation of the oligomerisation region of small terminase. Strikingly, for two protomers, density for the N-terminal residues Asp3-Val23 of one protomer and the C-terminus – residues Gly125 to Asp145 – of the adjacent protomer is clearly resolved in the map. The N-terminus of one protomer and the C-terminus of the adjacent protomer each form an α-helix that interact in a similar manner to the two helices in a classical Helix-turn-Helix (HTH) motif to create the DNA binding substructure. The orientation of these two helices is stabilised by 6 inter-helix hydrogen bonds **(Supplementary Fig. 10)** and the buried surface area between the helices is 430 Å^2^. These helices form a scaffold that places the N-terminal arm (NTA) of the N-terminal helix in the minor groove, while the end of the C-terminal helix is inserted into the major groove where the DNA is bent as it exits the central channel.

Although a 31bp oligo (7-37 of the *cos* sequence) was used to prepare the complex for cryoEM, only 28 base pairs are clearly resolved in the cryoEM density. The density for base pairs inside the channel is better resolved with density for nucleotides becoming less well defined as the DNA extends away from small terminase. Importantly, it was possible to unambiguously assign each nucleotide in the DNA within the channel. This enabled us to define the orientation of the dsDNA within the complex: the less well-ordered end that extends beyond the protein is the portion closest to the *cos* site. Therefore, in the complex the orientation of DNA is as follows: the *cos* cleavage site is positioned extending away from the surface of the small terminase where the DNA binding substructure is located, with the small terminase binding site bound partially within the tunnel and partially outside the nonamer by the DNA binding substructure. (**Fig. 3b**).

**Figure 3.**
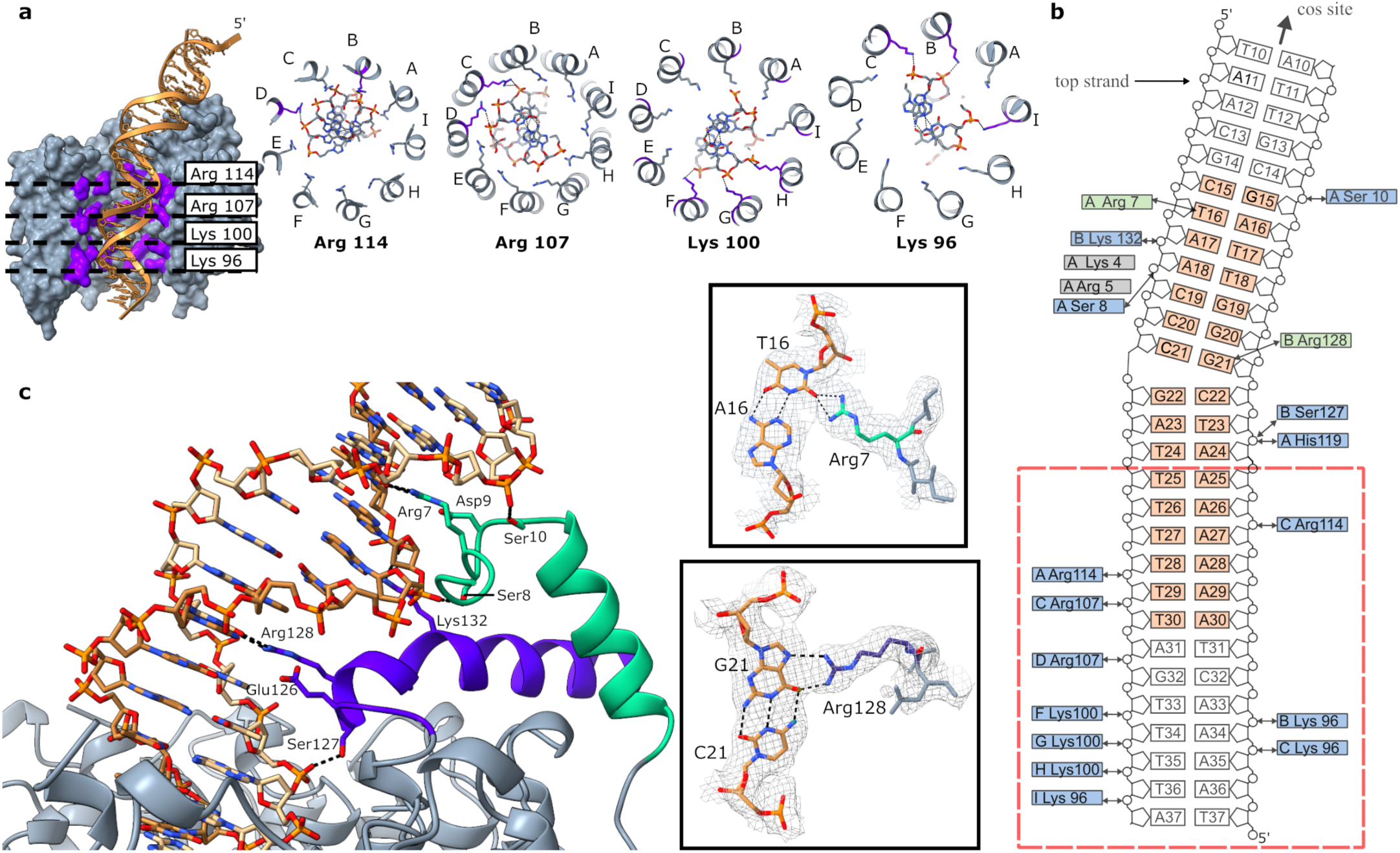
Interaction of small terminase with dsDNA. **a** Cross section of small terminase (grey) showing four positively charged rings formed by arginine and lysine residues (purple) with DNA depicted in gold. Protomers are labelled with letters A to I. **b** Schematic representation of overall small terminase: DNA interaction. The *cos* cleavage site is positioned 10 nucleotides upstream of the DNA shown. Residues forming specific interactions with DNA are shown in green and residues forming hydrogen bonds with the backbone phosphates of DNA are shown in blue. The interaction site on DNA for residues shown in grey was not unambiguously defined in the cryoEM density. The minimal small terminase binding site is shown in orange. Red dashed line outlines the position of the small terminase channel. Black arrows indicate hydrogen bonds. **c** The specific DNA binding substructure. CryoEM density along with corresponding models of Arg7:T16 and Arg128:G21 are shown in side boxes. Hydrogen bonds are shown by black dashed lines.

### Interactions with DNA bound in the central channel

The internal surface of the channel of small terminase is positively charged (**Fig. 2e, 3a**). Four rings of lysine and arginine residues: Lys96, Lys100, Arg107 and Arg114 line the channel and form hydrogen bonds with phosphates of the DNA backbone (**Fig. 3a**). The DNA duplex forms hydrogen bonds with 2-3 residues from adjacent protomers at each ‘level’, as it threads through the central channel. The composition of the group of protomers that contact the DNA at each level, rotate around the nonamer with respect to the DNA binding substructure. For example, the DNA forms bonds with three Lys96 (protomers B, C and I), followed by three Lys100 (from adjacent protomers F, G, and H) deeper in the tunnel, then Arg107 from two adjacent protomers (C and D) and finally Arg114 from two protomers (A and C) adjacent to the tunnel entrance closest to the DNA binding substructure formed from protomers A and B. His119 and Ser127 from protomers A and B, respectively, located at the top of the channel additionally form hydrogen bonds with the phosphates of the DNA (**Fig. 3b**). As DNA threads though the central channel, the interaction with the rings of charged residues lining the channel distort the DNA structure away from the canonical B-form resulting in narrowing of the minor groove and widening of the major groove (**Supplementary Fig. 11**). This coincides with the A-tract in the sequence of the minimal specific binding site.

### Specific protein: DNA interactions formed by the DNA binding substructure

Direct hydrogen-bonding interactions of small terminase with bases of DNA are formed solely by the DNA binding substructure. The two helices forming the DNA binding substructure position the sidechains of Arg128, which lies at the N-terminal end of the C-terminal helix, and Arg7 which is positioned in the NTA adjacent to the N-terminal helix, so they can reach into and contact bases in the major and minor grooves respectively. Arg128 forms bidentate hydrogen bonds with the base of G21 in the centre of the small terminase binding site. (**Fig. 3c**). Arg7 forms bifurcated hydrogen bonds with T16 at the 5′ end of the small terminase binding site. The position of both arginine sidechains is stabilised by ionic interactions with negatively charged residues Glu126 and Asp9 for Arg128 and Arg7, respectively. A similar mode of protein-DNA interaction was observed for the integration host factor interacting with the minor groove of DNA^25^. In both pairs of positively/negatively charged residues a serine residue (Ser127 and Ser8) lies between the residues, and its sidechain makes a hydrogen bond to the phosphate backbone of DNA (**Fig. 3c**). Additionally, in the NTA the sidechain of Ser10 makes a hydrogen bond to the other side of the major groove to the Ser8 contact, thus stabilising the interaction of the NTA with DNA that positions Arg7 in the minor groove. The distortion of the DNA upon binding, with the centre of the bend located where Arg128 is inserted into the major groove, is further stabilised by the interaction of Lys132 with the phosphate backbone on the opposite side of the major groove to Ser127. There are two more residues with positively charged sidechains in the NTA - Arg5 and Lys4. Whilst the density for the backbone in this region clearly shows that the NTA extends along the minor groove, there is no clear densisty to unambiguously define where these side chains contact DNA, suggesting the interaction could involve either contact to bases in the minor groove or the phosphate backbone.

### Influence of DNA sequence on binding

We investigated the importance of the nucleotide sequence of the binding site in the minor and major grooves **(Fig. 4)**. Swapping the orientation of the A:T base pairs in position 16 (where Arg7 contacts DNA) or position 18 (a possible interaction with Lys4) had little effect on DNA binding - as expected because A:T and T:A are indistinguishable in the minor groove^26^. In contrast, swapping the orientation of the C:G base pair in position 21 substantially reduced binding and substitution of the C:G base pair for a T:A base pair reduced DNA binding to a level undetectable by EMSA. This confirms that Arg128 in the C-terminal helix of the DNA binding motif plays a crucial role in specific protein:DNA interaction. The DNA is bent to an angle of 34° **(Fig. 1f)** at the point at which Arg128 forms hydrogen bonds with G21 **(Fig. 3c)**. The bending results in distortion of the DNA away from B-form with the minor groove widened and the major goove narrowed at the region centred on G21 (**Supplementary Fig. 11)**.

**Figure 4.**
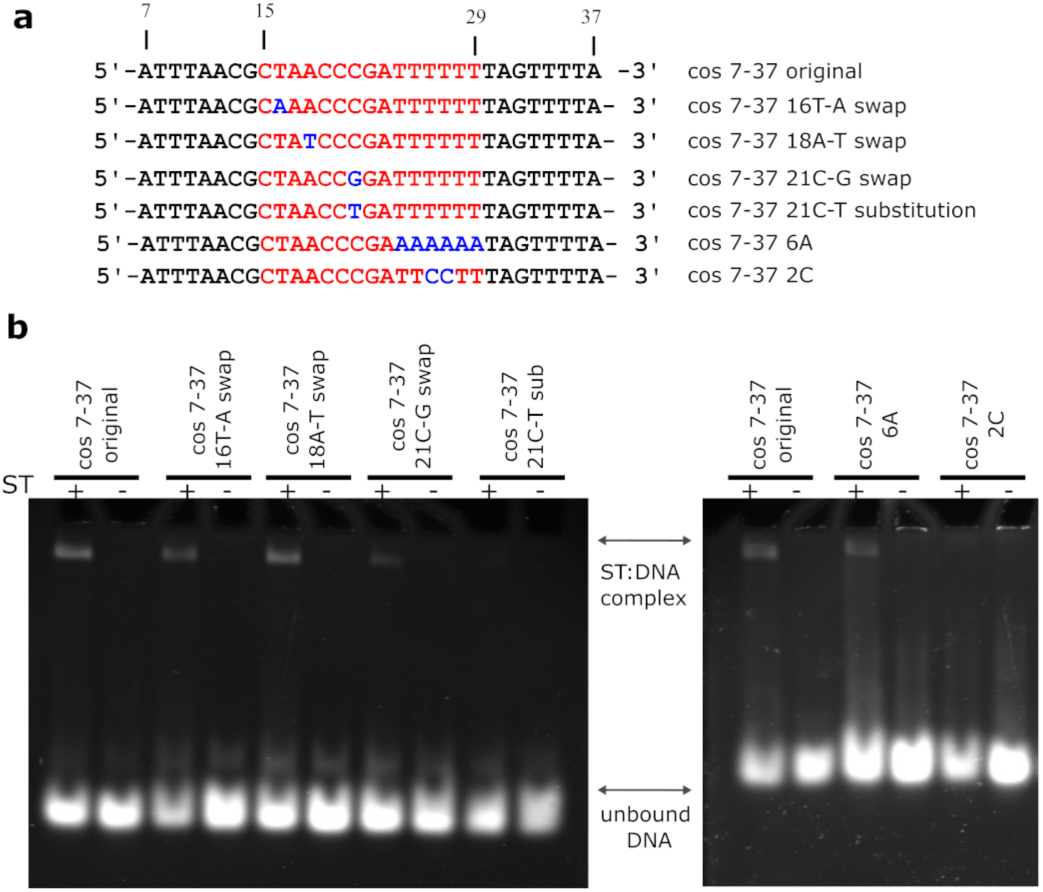
Interaction of small terminase with DNA mutants. **a** DNA sequences showing mutations in the small terminase binding site. Depicted in red is the region of DNA binding site. Mutations are highlighted in blue. **b** EMSA of wild-type and mutant DNA oligos with small terminase.

To further investigate the influence of the DNA sequence on the specificity of small terminase binding we designed two mutant DNA oligos, modifying the composition of the region of the binding site located in the central channel (**Fig. 4**). As would be expected given the non-sequence specific interaction between small terminase and DNA within this region, swapping the orientation of A:T base pairs at positions 24-29 had no influence on DNA binding. In contrast, introducing two GC base pairs at positions 26-27 in the middle of the A-tract abolished DNA binding. This indicates that while there are no base specific contacts within the tunnel, the flexibility of the overall DNA binding sequence, including the flanking regions, accorded by this A:T rich region, is essential for binding.

### Role of positively charged residues in interaction with DNA

To define the contribution of arginine and lysine residues in small terminase:DNA interaction we produced mutant proteins replacing several of these residues by alanine. Three residues within the central channel were mutated: Lys100, Arg107 and Arg114. Additionally Lys4, Arg5, and Arg7 in the NTA of the DNA binding substructure were individually mutated and a triple mutation of all three residues was also produced. Size exclusion chromatography (**Supplementary Fig. 12**) and circular dichroism (**Supplementary Fig. 13)** confirmed all mutant proteins were correctly folded and assembled into nonamers.

No DNA binding by any of the small terminase mutants could be detected by EMSA using a fluorescently labelled 100bp DNA duplex (**Fig. 5a**). Thus, to obtain information on binding affinity we performed more sensitive microscale thermophoresis (MST) experiments using the same DNA **(Fig. 5b, Supplementary Fig. 14)**. The equilibrium binding constant (K_d_) for DNA binding by WT small terminase was determined as 0.50 ± 0.03 μM (n=4). Either mutation in the central channel, Arg107Ala, or the mutation of 3 residues from the DNA binding motif, Lys4Ala/Arg5Ala/Arg7Ala, reduced the affinity by ∼1.5x: K_d_ = 0.79 ± 0.23 μM (n=3) and 0.85 ± 0.05 μM (n=6), respectively. To confirm the specificity of DNA interaction measured using MST, a 71bp DNA fragment missing the binding site, labelled with Cy5, was used in a control experiment. In this case the K_d_ for non-specific binding by WT small terminase was 2.04 ± 0.12 μM (n=3), four times lower affinity than for the 100bp DNA containing binding site.

**Figure 5.**
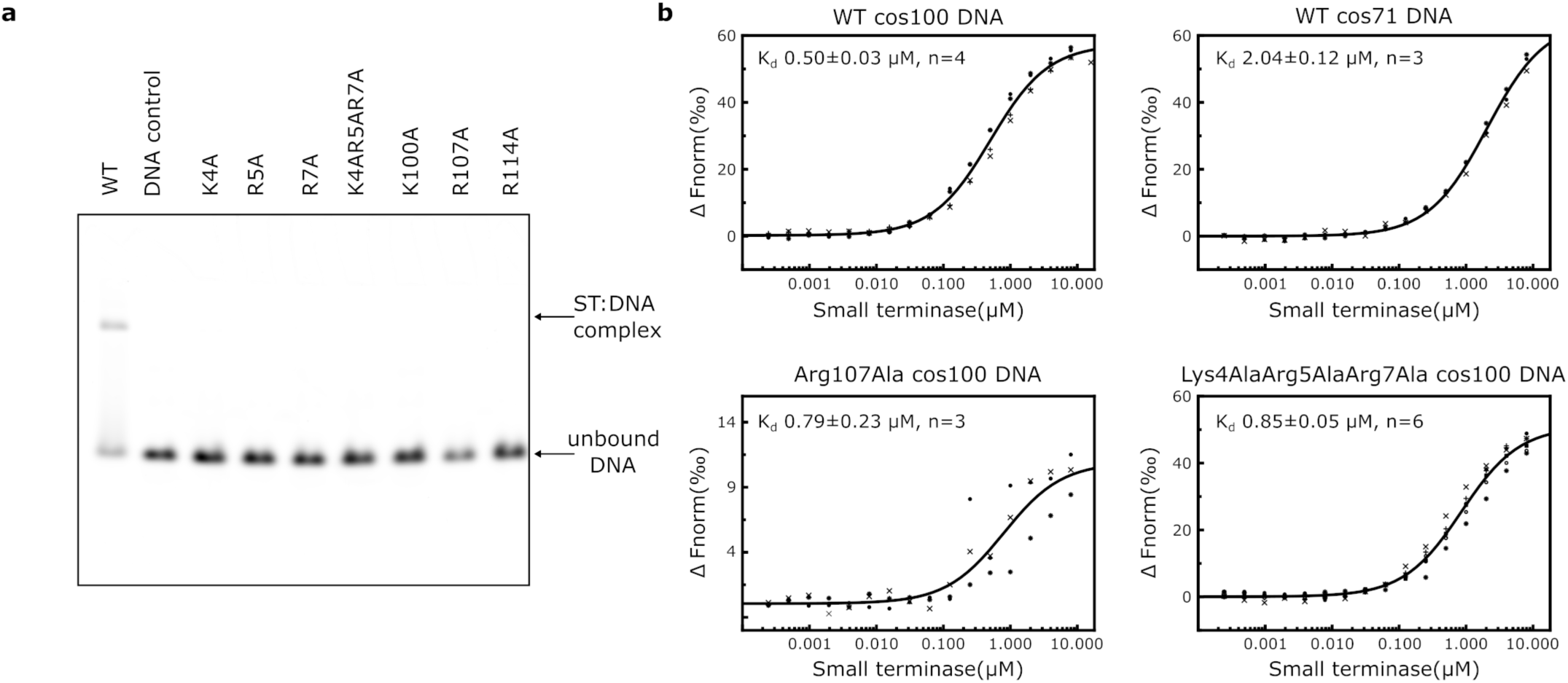
Interaction of small terminase mutants with DNA. **a** EMSA of small terminase (ST) mutant proteins with 100bp DNA labelled with Alexa 647 (cos100). **b** MST binding curves of the same dsDNA with WT and selected mutants and WT with DNA lacking the small terminase binding site (cos71). All measurements were repeated as 3 to 6 independent experiments.

## Discussion

### Comparison of small terminase structures of double stranded bacteriophages

We report the first experimentally derived structure for a bacteriophage small terminase protein bound to DNA, unveiling structural events that occur during recognition of viral genomic DNA by the small terminase. To date no common mechanism by which these proteins bind to DNA has been elucidated but several conflicting models have been proposed^9, 12, 14, 19^. One explanation for the wide variety of models that have been proposed is that for small terminase from several bacteriophages, the DBD can be produced as an isolated domain and structures of DBDs have been determined for small terminases of several phages including SF6 and λ^10, 11, 21^. Conversely, in small terminases from other bacteriophages the annotation of the putative DNA binding domain is less straightforward. Indeed, in some cases – P22, 44RR and HK97 – no well-defined DNA binding domain was known. The very N-terminus is well defined only in the structures of small terminase from G20c and λ phage. Intriguingly there is at least one positively charged residue at the N-terminus of all small terminases (**Supplementary Table 3**) suggesting that, as we observe for the HK97 small terminase, unstructured regions at the N-terminus of small terminase from other bacteriophages may also become ordered upon DNA binding. This is supported by data for the Sf6 small terminase, where it was shown that a Lys6Ala mutant in the unstructured N-terminal region abolished DNA binding^14^.

In all small terminase structures, the internal surface of the channel is positively charged (**Supplementary Fig. 1**). It is tempting to speculate that, as for HK97, DNA may bind in the central channel of other small terminases, but it is unlikely that this is the only mode of DNA interaction in all small terminases. Certainly, for small terminases from T4 and SF6, the assembled central oligomerisation domain showed significantly reduced DNA binding activity^9, 27^. In the case of T4 these experiments used EMSA and we note that this technique was also unable to detect binding by HK97 small terminase when mutations were made in the DNA binding substructure. However, tunnel mutants for T4 did show minimal effect on DNA binding^27^. Crucially for SF6, there is experimental evidence indicating that DNA does not interact with residues in the central channel^9^. The sizes of the central channels vary between small terminases from different bacteriophages (**Supplementary Fig. 15**) and measurement of the diameter suggests that whilst the channel of small terminase from G20c, P74-26, E217, 44RR and HK97 is wide enough to accommodate B-form DNA without any clashes, in small terminases from P22, SF6, SPP1, Sf6, PaP3 and pHBC6A51, the channel is too narrow for DNA to pass through.

In all small terminase structures determined to date, residues at the C-terminus are disordered, but the number of disorded residues ranges from very few (such as in SF6) to more than 40 residues (HK97, G20c, 44RR, E217). These unstructured regions are clearly visible as diffuse density in cryoEM 2D classes of E217^20^ and P74-26^18^. The only previous evidence that the C-terminus was involved in DNA binding was for P22 phage^17^ but given the contribution to DNA binding from the C-terminal region of HK97 small terminase where Arg128 makes a key sequence specific contact with DNA, it seems possible that the unstructured C-terminal regions of small terminase from other bacteriophages may also become ordered on DNA binding.

### HK97 small terminase recognises DNA using a unique ‘arginine clamp’

To our knowledge, the structure of HK97 small terminase bound to DNA reveals a hitherto unknown mode of DNA binding. Our data suggest that both the bending of DNA and the interaction with Arg128 are crucial for sequence specific DNA recognition by HK97 small terminase because mutation of guanidine 21 abolishes DNA binding, while swapping DNA bases in the minor groove had a minor influence on interaction (**Fig. 4)**. The positioning of 2 arginine residues into the major and minor grooves of the DNA form an ‘arginine clamp’ that locks the bent conformation of DNA into place. Further residues from the DNA binding substructure hydrogen bond to the phosphate backbone and stabilise the clamp’s grip on DNA. This interaction relies on the distortion to the DNA, facilitated by the run of A:T base pairs^26, 28^ within the central channel adjacent to the bend, since disrupting the propensity for bending of this sequence by the inclusion of C:G base pairs abrogated small terminase binding **(Fig. 4)**. The clamp ensures that small terminase tightly binds to its binding site.

### A model for packaging termination in HK97 phage

The small terminase is essential for termination of packaging at the *cos* sequence^4, 5^, and data presented here enable us to suggest a model of packaging termination in *cos* bacteriophages (**Fig. 6**). We propose that during DNA packaging the small terminase encircles the DNA. Available evidence enables us to propose the following mechanism for packaging termination. i) while the large terminase mechanically translocates DNA into the capsid, the suggested conformational changes in all five terminase subunits propagating around the ring^23^, would result in dislocation of small terminase. This continuous displacement of small terminase, coupled with the path DNA takes as it threads through the channel, suggests that as small terminase slides along the genomic DNA it may rotate with respect to large terminase. During this process the flexible C-terminal regions of small terminase surve as molecular sensors surveilling the DNA exiting from the central channel. ii) as the 9bp long A-tract begins to exit the channel, the flexibility of the junction between this and the adjoining C:G base pairs, allows the DNA to bend. Shape matching of the bent DNA with the C-terminal region of one protomer positions its α-helix so that the side chain of Arg128 can insert into the major groove, specifically interacting with G21. iii) the N-terminal region of the adjacent protomer then folds into an α-helix that stacks alongside the α-helix from the C-terminus to complete the DNA binding substructure and the NTA inserts into the minor groove with Arg7 making contact with T16. This closes the ‘arginine clamp’ which locks small terminase onto DNA. iv) the bent conformation of the bound DNA results in an interaction between small and large terminase in which the central axes of the circular assemblies of the small and large terminase oligomers no longer align, which stalls DNA translocation and v) triggers the switch of large terminase from DNA translocation to nuclease activity. Stalling of translocation in HK97 has been observed *in vitro* at experimentally-induced low packaging velocities^4, 23^. The specific recognition of G21 by Arg128 and the adjacent A-tract via the shape of the DNA ensure that stalling and subsequent triggering of nuclease activity occur only when the *cos* cleavage site is accommodated at the nuclease active site of the large terminase. The 10 nucleotide overhang at the *cos* cleavage site may be a result of an initial contact with small terminase triggering the cut of the first strand and subsequent DNA translocation as the ‘arginine clamp’ is released from the binding site before the second strand is cut. Finally, the terminase complex dissociates from the capsid vertex ready for the next packaging event.

**Figure 6.**
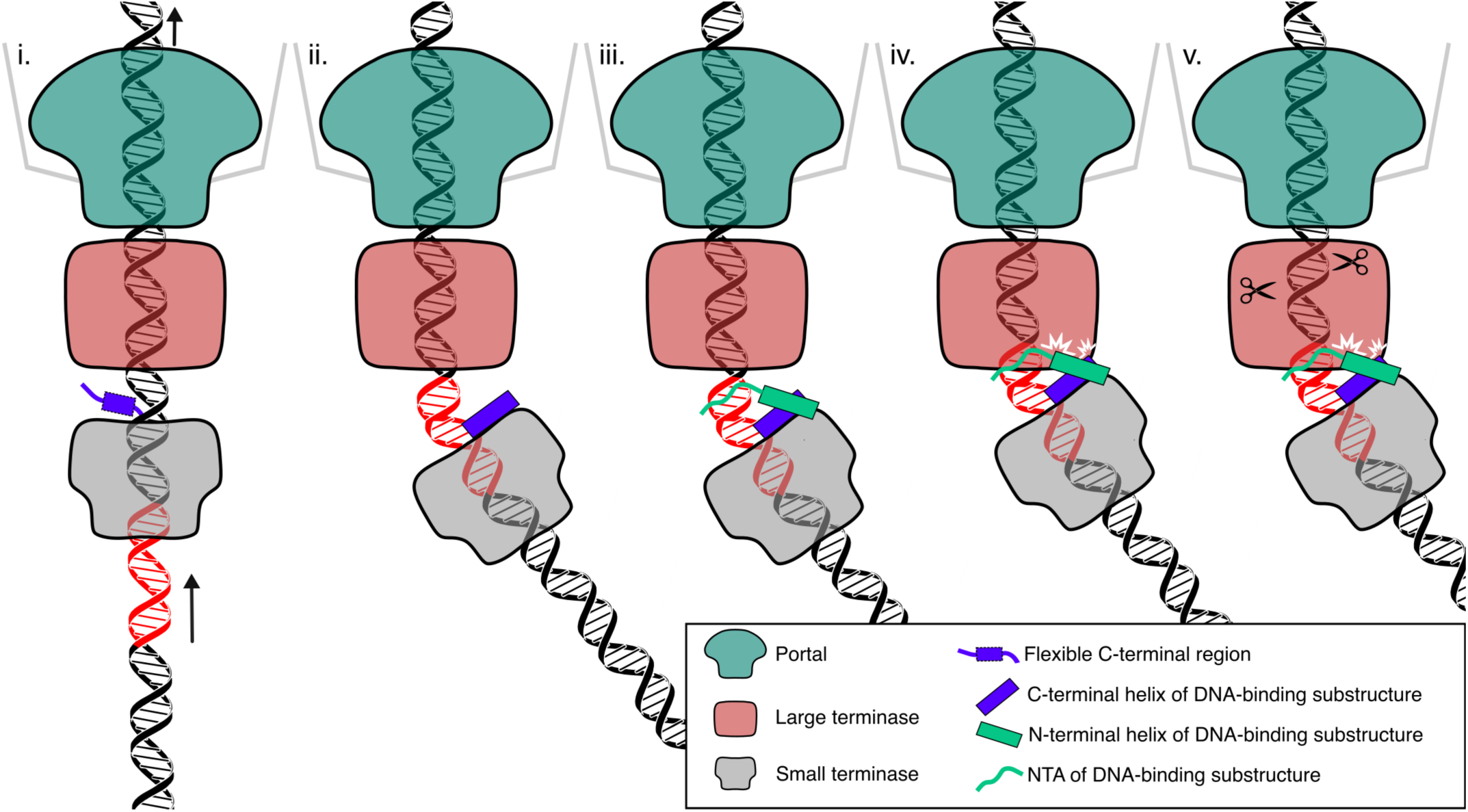
Proposed mechanism of packaging termination. Roman numerals refer to stages described in the text. Portal protein 12-mer, large terminase 5-mer, small terminase 9-mer, NTA and helices forming the DNA binding substructure and flexible C-terminal segments indicated in legend. The small terminase binding site is highlighted in red on the DNA. Arrows indicate the direction of the DNA movement.

## Conclusions

In this work we defined the small terminase binding site in the HK97 genome which enabled us to derive the first structural data for a viral small terminase protein in complex with DNA. The structure reveals a hitherto unknown mode of DNA recognition by a circular protein that can slide along DNA. Upon encountering the specific DNA binding sequence, two disordered regions at the N-terminus and C-terminus, respectively, of two adjacent protomers, fold into α-helices and form a DNA binding substructure that locks small terminase onto DNA. The unique properties of this protein allowing threading along DNA while transiently surveying its sequence until a specific site is reached and arrested by the protein, make it a valuable tool for potential biotechnological applications. The structural findings, unveiling molecular events that occur during viral DNA recognition by the *cos* bacteriophage HK97 may be applicable for understanding how other systems, involving circular proteins that bind to DNA, work.

## Methods

### Cloning

The gene encoding HK97 small terminase (residues 1-161) was cloned in Champion pET SUMO vector (ThermoFisher)^24^. Mutants of small terminase were produced by site directed mutagenesis with the original plasmid as a template and a pair of oligos (Eurofins Genomics) containing mutated region.

### Protein Expression and Purification

Small terminase was overexpressed in *E*. *coli* BL21(DE3) Gold pLysSRARE cells. Cells were grown in 1L LB media supplemented with 30 µg/mL kanamycin and 34 µg/mL chloramphenicol at 37°C till OD_600_ reached 0.6. Then the flasks were cooled on ice and expression was induced with 0.5 mM IPTG. Cells were incubated at 16°C overnight and harvested by centrifugation. The cell pellet was frozen at -80°C. Cells were resuspended and lysed by sonication in buffer A: 50 mM Tris-HCl, pH 7.5, 1M NaCl, 5% (v/v) glycerol, 30 mM imidazole supplemented with 100 µM AEBSF (4-(2-aminoethyl) benzenesulfonyl fluoride), 0.5 µM leupeptin, 0.7 µM pepstatin, 100 µg/mL lysozyme, 10 µg/mL RNaseA, 2.5 µg/mL DNase, 1mM MgCl_2_. After centrifugation for 1 hour at 18K rpm, cell lysate was applied to HisTrap FF Ni column (Cytiva). To reduce nucleic acid contamination, the column was washed with 10 column volumes (CV) of buffer A (resuspension buffer without protease inhibitors and other additives), followed by 5 CV of high salt buffer (25 mM Tris-HCl, pH 7.5, 3M NaCl). After re-equilibration of the column with buffer A, the protein was eluted with 50 mM Tris-HCl, pH 7.5, 1M NaCl, 5% (v/v) glycerol, 0.5 M imidazole. Eluted protein was diluted twice with 25 mM Tris-HCl, pH 7.5 to reduce NaCl concentration to 0.5 M and incubated with 1:100 (w/w) SUMO protease overnight at 4°C with addition of 2 mM DTT. Cleaved protein was diluted further to 0.2M NaCl concentration and applied to cation exchange 10/10 Mono S column (GE Healthcare). Small terminase was eluted in two peaks with 0.2- 1M NaCl gradient. Fractions from each peak were concentrated separately in Vivaspin concentrators (Sartorius) with MWCO 30K cut off and loaded onto a Superdex 200 10/300 column (GE Healthcare) in 25 mM Tris-HCl, pH 7.5, 1M NaCl buffer. Finally, a Heparin column (GE Healthcare) was used for the polishing step. Protein was loaded on the column in 25 mM Tris-HCl, pH 7.5, 0.15 M NaCl buffer and was eluted with a linear gradient 0.15-1 M NaCl. Protein was concentrated and flash frozen in liquid nitrogen and stored at -80°C.

Small terminase mutants were expressed in the same way as WT protein with the following modifications. For Ni affinity purification a batch binding method was used. Cleared cell lysates were mixed with 5 mL Ni-NTA affinity resin (Generon) equilibrated with buffer A and incubated by rotating the tubes for 30 min at room temperature (RT). Resin was poured into 10 mL empty PD-10 columns (GE Healthcare) and all washes and elution were performed by gravity flow with buffers as used for WT purification. Additionally, a heparin column was used after SUMO protease cleavage instead of a cation exchange column. This step helped to separate cleaved protein from SUMO tag as well as minimize nucleic acid contamination. Finally, all mutants were further purified by size exclusion chromatography on a Superdex S200 10/30 column in 25 mM Tris-HCl, pH 7.5, 1M NaCl, concentrated and frozen in liquid nitrogen.

### Preparation of DNA oligos

Fluorescent DNA fragments of 100 bp (*cos*-20-80) and 71 bp (*cos*31-100) were produced by PCR reaction with DreamTaq DNA Polymerase (ThermoFisher Scientific). The forward primer (Eurofin Genomics) was covalently linked with Alexa 647 for 100 bp DNA and the reverse primer with Cy5 for 71 bp DNA. PCR settings: initial cycle 94 °C - 30 s followed by 35 cycles of 94 °C – 10 s, 60 °C – 30 s and final extension of 72 °C – 5 min. A plasmid containing 784bp segment around the *cos* site (-312 +472) was used as template in all reactions at a final concentration of 2 pg/µL^24^. The final PCR products were purified using a PCR clean-up kit (Macherey-Nagel).

Short oligos for EMSA and complex formation (Sigma-Aldrich) were reconstituted with TE buffer (10 mM Tris-HCl, pH 8.0, 1 mM EDTA) to a final concentration of 200 µM. Equimolar quantities of complementary oligos were mixed in a PCR tube and annealed in a heating block (5 min at 95 °C followed by slow cooling over 2 h to RT). The quality of annealing was checked on 10% acrylamide native gel. Annealed oligos were diluted with water to 10 µM before use.

### Electrophoretic mobility shift assay (EMSA)

For each native gel a master mix (MM) was prepared containing small terminase in 25mM Tris, pH 7.5, 200 mM NaCl, 10 mM MgCl_2_ buffer. 8 µL of MM were aliquoted into each tube and then 2 µL of 10 µM DNA was added followed by thorough mixing. Final concentrations of small terminase and DNA in 10 µL reaction were 1 µM and 2 µM respectively. Tubes were incubated at RT for 3 h. 3 µL of loading buffer (50% glycerol, Orange G) was added to each tube, mixed, and spun for 1 min at 13K rpm before loading on 6% native gel. A gel with acrylamide:bis-acrylamide 99:1 ratio was prepared using running buffer 25 mM Tris-HCl pH 8.3, 192 mM Glycine, 200 mM Na_2_SO_4_, 10 mM MgSO_4_ and poured into empty 1.5 mm thick gel cassettes (ThermoFisher Scientific), which made handling of low percentage gels easier. Gels were run overnight at 4 °C with current of 15 mA to prevent overheating. Gels were rinsed with TAE buffer followed by staining with 0.2% EtBr in TAE buffer for 30 min and imaged with GelDocXR+ (BioRad).

30% acrylamide:bis-acrylamide (99:1 ratio) solution was prepared by mixing 7.4 mL of 40% acrylamide (BioRad), 1.5 mL of 2% Bis-Acrylamide (Biorad) and 1.1 mL of water. Such acrylamide:bis-acrylamade solution was used to prepare gels with a larger pore size to facilitate movement of the large protein:DNA complex through the gel.

EMSA for small terminase mutants was performed as described above. 100 bp fluorescent oligo labelled with Alexa647 containing *cos* site (*cos* -20-80), was used at a final concentration of 30 nM. MM containing fluorescent oligos in 25 mM Tris, pH 7.5, 200 mM NaCl, 10 mM MgCl_2_ was aliquoted into the tubes, protein was added to a final concentration of 1 µM, mixed and incubated for 3 h at RT. Gels were run at 4 °C overnight as described above and scanned with GE Amersham Typhoon-5 fluorescence gel and blot scanner (GE Healthcare) with red laser.

### Far-UV circular dichroism spectro polarimetry (CD)

Proteins (WT and mutants) were diluted to a final concentration of 0.3 mg/mL with 25 mM Tris-HCl, pH 7.5, 100 mM Na_2_SO_4_ in a total volume of 400 µL. Spectra were collected on a Jasco J-1500 Circular Dichroism Spectrometer from 260 nm to 195 nm with 1 mm pathlength.

### Microscale thermophoresis (MST)

Protein and DNA were diluted in reaction buffer: 25 mM Tris-HCl, pH 7.5, 200 mM NaCl, 10 MgCl_2_, supplemented with 0.005-0.01% Tween 20 (NanoTempo Technologies). To minimize pipetting errors 1:1 serial dilutions were performed such that protein was diluted in 12 nM DNA solution. For each MST run 2x stocks of small terminase and fluorescent DNA oligos were prepared at 16 µM and 24 nM respectably. Stocks were spun for 3 min. DNA was diluted twice with reaction buffer to 12 nM and then 10 µl was aliquoted in 15 tubes. 10 µl of each 2x stock was mixed in a separate PCR tube for the highest protein concentration followed by serial dilution in 15 consecutive tubes. Tubes were spun again for 5 min at 13K rpm before loading in Premium Monolith NT.115 Capillaries (NanoTempo Technologies). This produced a series of 16 samples with small terminase concentration ranging from 0.24 nM to 8 µM and constant DNA concentration of 12 nM. For one run a 2x stock of WT small terminase at 32 µM was used resulting in final protein concentrations ranging from 0.48 nM to 16 µM. MST was measured using a Monolith NT.115 instrument (NanoTemper Technologies) at an ambient temperature of 20 °C. Instrument parameters were adjusted to 60 % LED power and medium MST power (40%). Data from three or more independently pipetted measurements were analyzed (MO.Affinity Analysis software version 2.3, NanoTemper Technologies) using the signal from an MST-on time of 5 s. Fnorm data were fit to a model describing a molecular interaction with a 1:1 stoichiometry according to the law of mass action, to determine the equilibrium binding constant (K_d_), where fraction of DNA bound f(c), at a given protein concentration (c) for a constant total DNA concentration (c_T_) in the assay is defined as:

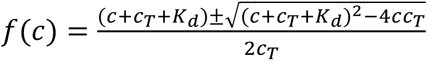. This model is adjusted to account for normalized fluorescence (Fnorm) of bound and unbound DNA as follows: *fnorm = (1 - f(c))unbound + f(c) bound*. Individual datapoints for each set of experiments were globally fit to this model using the Levenberg-Marquardt Algorithm in pro Fit 7 (Quantumsoft), allowing all parameters to float globally except for the DNA concentration (c_T_), which was held at 12 nM for all datasets. Data and fits were plotted as ΔFnorm. Residuals for each fit (datapoint-fitvalue) were calculated for each datapoint used in the fit and plotted as Fraction ΔFnorm (**Source Data**). K_d_s are listed with ± standard deviation.

### Small terminase:DNA complex formation and cryoEM sample preparation

Small terminase:DNA complex was prepared at 1:2 protein to DNA ratio by mixing small terminase with 31 bp DNA (*cos* 7-37) in reaction buffer 25mM Tris-HCl, pH 7.5, 200mM NaCl, 10mM MgCl_2_. After incubation for 3 hr at RT, it was further diluted with reaction buffer to 400 µL and final concentration of small terminase and DNA 11.9 µM and 24 µM respectably. The small terminase:DNA mixture was loaded onto a S200 10/30 column (GE Healthcare) equilibrated with reaction buffer and run at 0.5 ml/min. 100 µL fractions were collected. The most concentrated fraction was spun for 5 min at 13K rpm before applying 3 µL to UltraAuFoil R1.2/1.3 gold support grids (Quantifoil). Prior to sample applications grids were glow-discharged for 3 min in Pelco easiGlow glow-discharger (Pelco) at 20 mA, 0.38 mBar. Grids were blotted at 4°C and 100% relative humidity for 2 s with -20 blot force and vitrified by plunging into liquid ethane using the FEI Vitrobot Mark IV (Thermo Fisher). Grids were prepared within one hour of completion of the gel filtration run. Complex formation was later confirmed by running fractions on native acrylamide gel.

### CryoEM data acquisition

Initial screening of grids was performed in Sheffield on Tecnai Arctica (FEI) 200kV cryo-electron microscope. Two separate data collections were done from the same grid using a Titan Krios microscope (FEI) operated at 300 keV and equipped with a Gatan K2-Summit detector and energy filter (a slit width of 20 eV was used) at Astbury Biostructure Laboratory, University of Leeds. Movies were collected automatically with EPU (Thermo Fisher Scientific) software on Gatan K2-Summit detector operated in counting mode with calibrated pixel size of 1.07 Å. For the 1^st^ dataset 682 movies were collected with target defocus range of 1.3 to 3.1 μm. Each movie comprised 50 frames with a total fluence of 53 e-/Å^2^ over 8 s, corresponding to a flux of 7.58 e-/pixel/s. For the 2^nd^ data collection 2085 movies were collected over a target defocus range of 1.0 to 2.4 μm with a total fluence of 54 e-/Å^2^ over 8 s, corresponding to a flux of 7.65 e-/pixel/s. To address any preferential orientation problem 995 movies were collected at 10° stage tilt and 71 at 20° stage tilt.

### Cryo-EM data processing (Supplementary Fig. 5-7, Supplementary Table 1)

Both datasets were processed in RELION 3.1.2^29^. Micrographs from the first dataset were motion-corrected using RELION’s implementation of the MotionCor2 algorithm^30^ . After estimation of contrast transfer function (CTF) parameters with CTFFIND-4.1^31^ micrographs were manually sorted to eliminate empty and icy ones. Micrographs collected at defocus greater than 3.8 µm were also discarded, resulting in 550 micrographs. Particle picking was performed with Topaz software^32^. Initially, 2700 particles were picked manually from a subset of 140 micrographs (denoised using Topaz). Topaz was then trained using these particle coordinates. The trained model was used to pick 250798 particles from the complete dataset. Particles were extracted in 168 pixel box and downsampled to 2.50 Å/pixel. 2D classification of selected particles showed predominately side views, although there were some classes with possibly top/bottom views (**Supplementary Fig. 5**). After 2D classification the best classes (250549 particles) were selected for 3D refinement. A map generated from the crystal structure of small terminase (PDB: 6z6e)^4^ low-pass filtered to 10 Å was used as a reference. A mask around the map from the crystal structure was used to focus the refinement on small terminase. The resulting map was used for partial signal subtraction in order to improve the signal for focused classification of the DNA region. We created a funnel mask in Chimera^33^ that masked DNA in the tunnel of small terminase as well as DNA above it.

3D classification focused on the region within this mask with no angular sampling and high T (50) produced classes with different DNA orientations. The most prominent class with 38582 particles was selected. Final resolution of the map after reverting subtraction, reconstruction, refinement and postprocessing was 5.0 Å. Re-extracting particles in 224 pixel box and downsampling to 1.25 Å/pixel, followed by CTF refinement and Bayesian particle polishing improved map resolution to 3.5 Å.

Movies from the second dataset were motion-corrected and CTF parameters estimated as described for the 1^st^ dataset (**Supplementary Fig. 6**). All icy and empty micrographs were discarded and additionally, micrographs collected at greater than 3 µm defocus or with estimated astigmatism greater than 600 Å were removed, resulting in 1874 micrographs. Particles were picked using the same Topaz model as used for 1^st^ dataset. 683470 particles were extracted from those micrographs in 168 pixel box and downsampled to 2.50 Å/pixel. A round of 2D classification was used to clean the dataset, resulting in 669682 particles. 3D classification was carried out with 7 classes using the final model from the 1^st^ dataset initially low pass filtered to 25 Å as the reference. Particles from the best class were re-extracted in a 224 pixel box and downsampled to 1.25 Å/pixel, 3D refined again and then CTF refined. Bayesian polishing was performed followed by another round of refinement. A final resolution of 3.9 Å was achieved. A second round of Topaz training was performed with selected particle coordinates from the final set of particles in the 2^nd^ dataset (**Supplementary Fig. 6**). Firstly, a subset of 517 micrographs containing more than 100 particles per micrograph was selected. 2D classification was run on 62616 particles from these micrographs and 250 random particles from 25 classes showing a variety of views were selected to create the set of 6250 particle coordinates used for Topaz training. From this point the workflow for both datasets were identical (**Supplementary Fig. 7**). Particle picking was performed with the new Topaz model from 550 and 1874 micrographs for the 1^st^ and the 2^nd^ datasets respectively. 306866 (dataset 1) and 837519 (dataset 2) particles were picked and extracted in 168 pixel boxes and downsampled to 2.50 Å/pixel. These particles were cleaned by 2D classification followed by 3D classification. Particles form the best classes were re-extracted in 224 pixel boxes and downsampled to 1.25 Å/pixel and 3D refined again, followed by Bayesian polishing with shiny particles extracted from movie frames in 320 pixel boxes and downsampled to 1.19 Å/pixel (final box size 288 pixels). After 3D refinement of the shiny particles the resolution of the maps was 3.1 Å and 3.2 Å for the 1^st^ and 2^nd^ dataset respectively. While performing Bayesian polishing, we noticed that particle movement graphs for each datasets looked very different. During the second data collection a number of movies were collected with the stage tilted at 10° or 20°. Eventually we discarded all tilted micrographs from this dataset. 3D refinement of 334433 particles from combining both datasets was followed by another round of CTF refinement and 3D refinement leading to an overall resolution of 2.9 Å. The final set of 334433 particles was split into 2 subsets: 174040 particles from micrographs collected close to focus (< 2 µm) and 160165 particles from micrographs collected further from focus (> 2 µm). The subset of particles from micrographs collected close to focus refined to identical overall resolution as the whole data set. To improve the density for the N-terminal helix that contacts DNA, we performed focused 3D classification on the close-to-focus subset, splitting particles in two classes with no angular sampling and high T (32) with partial signal subtraction using a mask containing the region of 2 helices and a segment of bound DNA (**Supplementary Fig. 7**). 3D refinement of 76626 particles from the best class resulted in slightly lower overall resolution of 3.0 Å, but better resolved density for the N-terminal helices.

### Model building and refinement (Supplementary Table 2)

Atomic model building in the final map was performed in Coot^34^ using crystal structure of HK97 small terminase (pdb 6z6e) as the initial model. Each chain was fit individually with jiggle fit option. Ideal poly-alanine α-helices were fitted into the extra density and then side chains were built. Two segments of ideal double stranded B-form DNA were fitted into DNA density and then joined together. It was possible to unambiguously assign the DNA sequence due to the high quality of the density for the middle region of DNA. Overall, 28 out of 31 DNA base pairs were modelled. No density was observed for residues 1-23 and 124-161 in 7 of the 9 protein chains. Real-space refinement was carried out in ISOLDE^35^ followed by reciprocal space refinement with REFMAC5/Servalcat^36^ using DNA restraints generated with libg in CCP-EM^37^ and cycles of manual re-building in Coot^34^. Model validation was done in CCP-EM by Validation:model software. All figures were generated by ChimeraX^38^ and Chimera^33^. To calculate the electrostatic potential, the PDB format files were converted to PQR format with the PDB2PQR server using the PARSE force field and assigned protonation states at pH 7.5. The APBS server^39^ was used and 0.2 M of ions were included in the calculation. Surfaces of small terminases in **Supplementary Fig. 1** were coloured by default electrostatic potential algorithm in ChimeraX. Hydrogen bonds were estimated with PISA server^40^. The angle of the DNA bend in Figure 6 was calculated between the vectors obtaining by aligning 10bp fragments of ideal B-form DNA with DNA from the structure at the bottom and the top termini.

## Data availability

Atomic coordinates and cryoEM maps have been deposited with the Protein Data Bank and Electron Microscopy Data Bank under accession numbers 8POP and EMD-17794 (map after focused classification), EMD-17818 (consensus map), respectively.

## Acknowledgements

This research was funded by the Wellcome Trust [206377 to AAA]. The authors would like to thank A. Leech and K. Hodgkinson for support and access to the University of York Bioscience Technology Facility, J. Turkenburg and S. Hart for help and support in using Tecnai12 microscope, S. Tzokov (University of Sheffield) for the opportunity to screen cryo-EM grids. CryoEM data were collected at the Astbury Biostructure Laboratory and we thank E. Hesketh and R. Thompson for assistance with data collection. We are also grateful for computational support received from the University of York High Performance Computing service (Viking). This project was initiated while SJG was employed at the University of York and is in no way related to her current role at Dreampore. Molecular graphics and analyses performed with UCSF ChimeraX, developed by the Resource for Biocomputing, Visualization, and Informatics at the University of California, San Francisco, with support from National Institutes of Health R01-GM129325 and the Office of Cyber Infrastructure and Computational Biology, National Institute of Allergy and Infectious Diseases.

## Author Contributions

MC, SJG and HTJ conceptualized the project and designed the research. AAA supervised the research. MC produced samples and prepared cryoEM grids, MC and HTJ collected cryoEM data and determined structures. MC performed binding experiments. MC, SJG and HTJ analysed data. MC, SGJ, HTJ and AAA wrote the manuscript.

