## Supplementary information for "Structure of HK97 small terminase:DNA complex unveils a novel DNA binding mechanism by a circular protein"

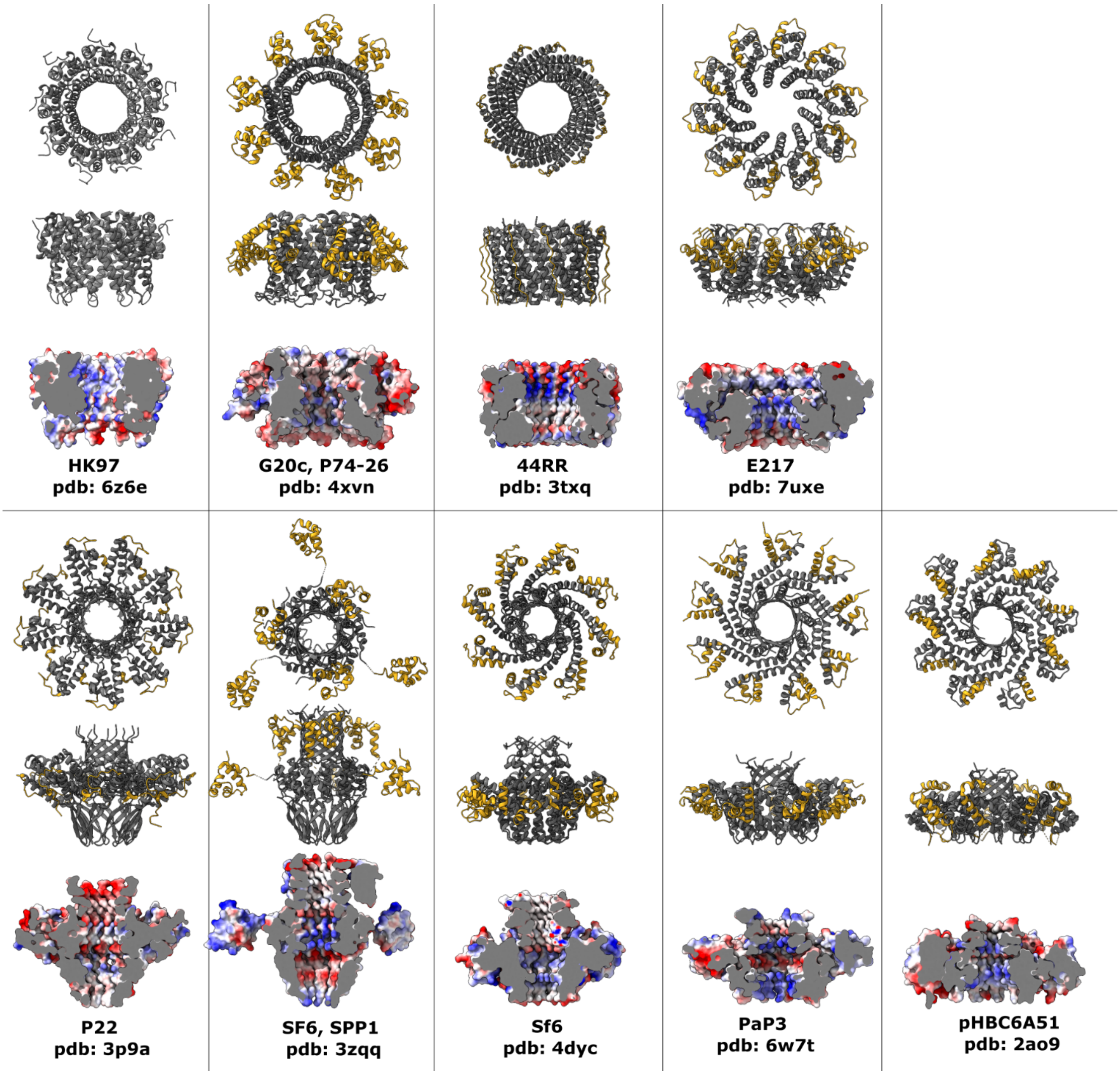

**Supplementary Figure 1. Structures of small terminases deposited in PDB and EMDB.** All structures are shown at the same scale: top and side view are presented with oligomerisation domain and C-terminus in grey and DNA binding domain (confirmed experimentally or predicted) in gold. Below is the cross section of the same structure coloured by electrostatic potential (red - negatively charged, blue - positively charged, range -16.7 to 16.7 kT/e).

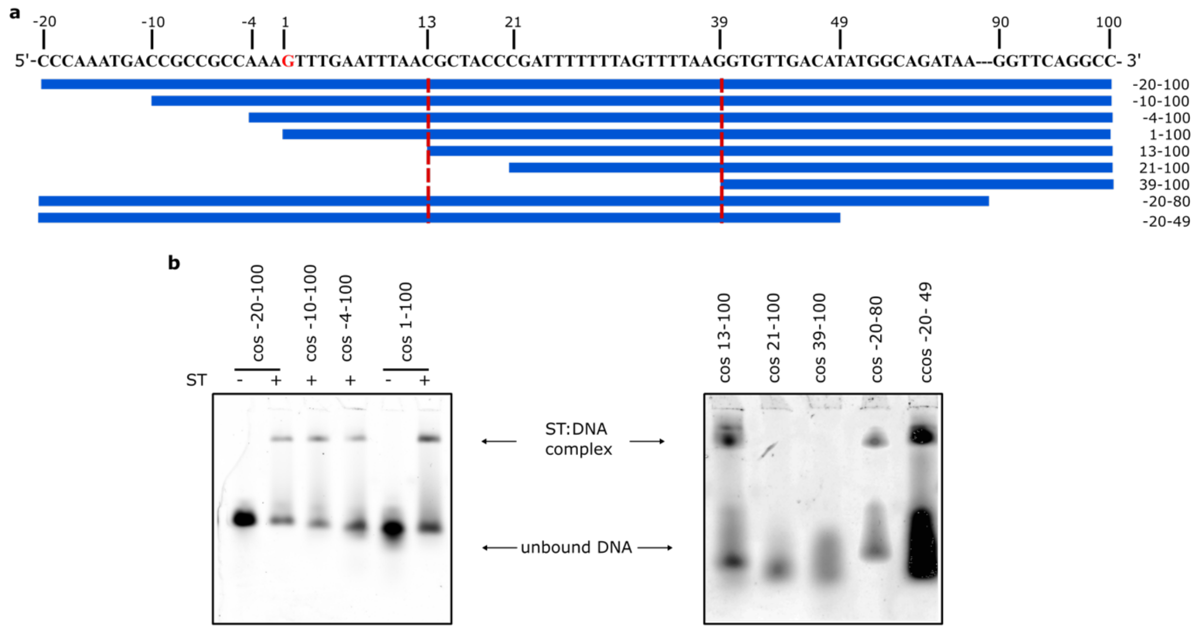

**Supplementary Figure 2. Determination of small terminase binding site. a** Sequence of HK97 putative small terminase binding site showing oligos used. Position 1 - cleavage site during genome packaging. The minimal small terminase binding site is marked with red dotted lines. **b** EMSA of interaction of fluorescent oligos spanning different regions of the putative binding site with small terminase.

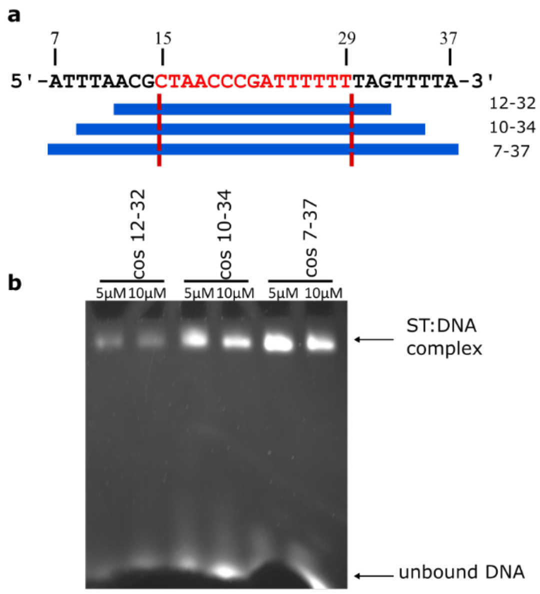

**Supplementary Figure 3. Oligo optimisation for small terminase:DNA complex formation. a** Sequence of small terminase binding site showing oligos used. **b** EMSA of the interaction of these oligos with small terminase.

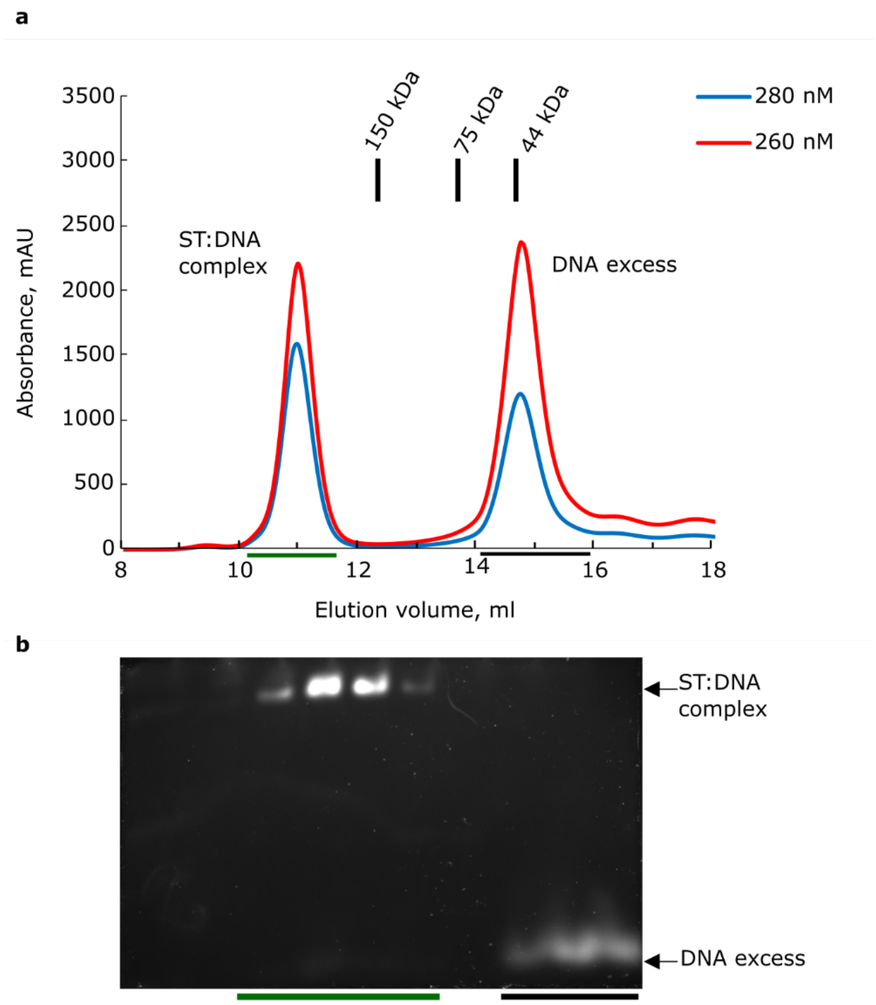

**Supplementary Figure 4. Purification of small terminase:DNA complex on S200 10/30 column. a** Analytical size exclusion chromatography profile. **b** Native SDS gel stained with EtBr, with elution fractions for the complex (green bar) and excess of DNA (black bar) labelled.

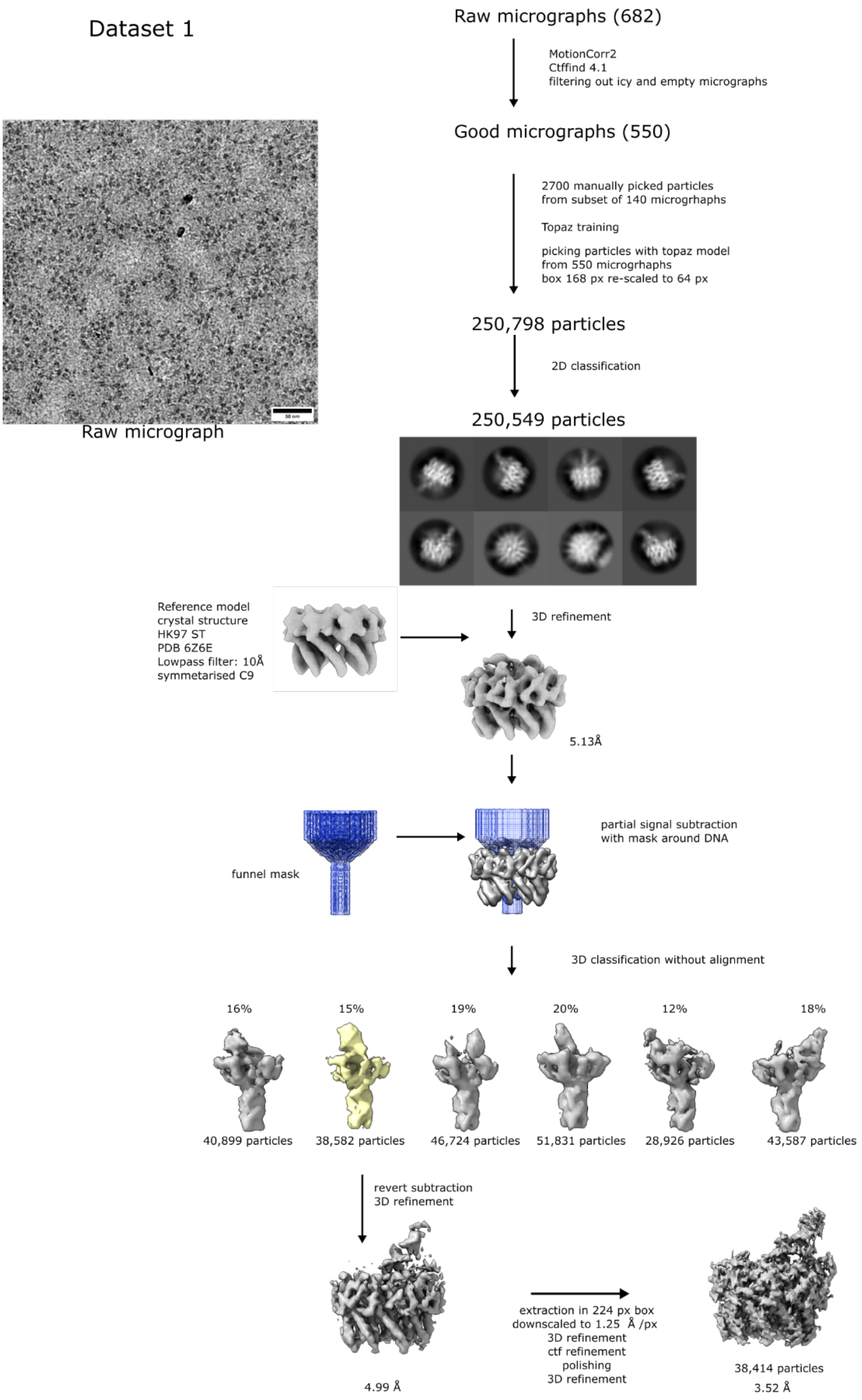

**Supplementary Figure 5**. **CryoEM processing flow chart for dataset 1.** Overview of initial cryoEM processing steps for small terminase:DNA complex dataset 1. Mask used in focused classification with partial signal subtraction is shown in blue. A representative micrograph showing particle distribution (scale bar is 50 nm) and a selection of 2D class averages representing different particle views is shown.

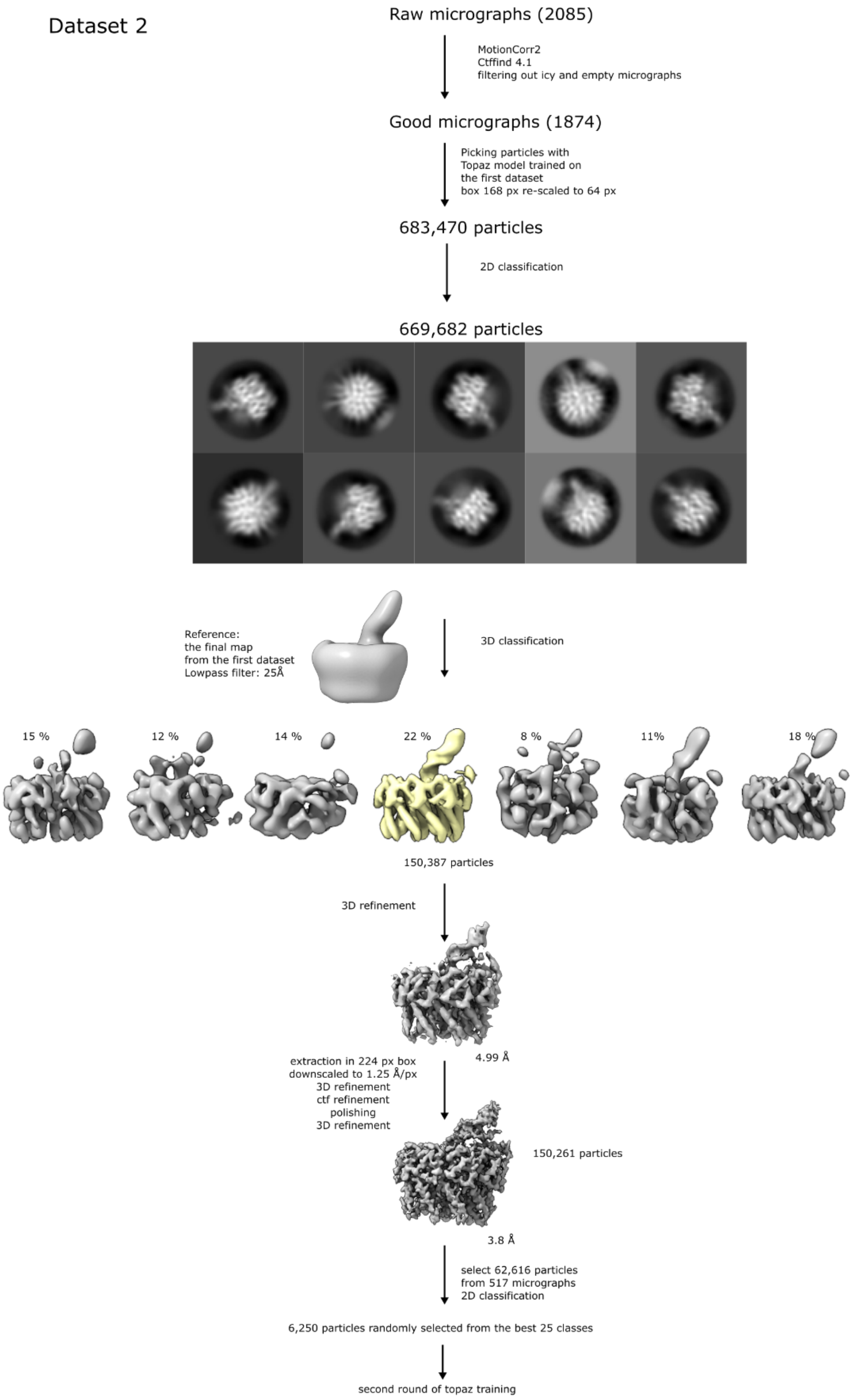

**Supplementary Figure 6**. **CryoEM processing flow chart for dataset 2.** Overview of the initial cryoEM processing steps for small terminase:DNA complex dataset 2. A selection of 2D class averages representing different particle views is shown.

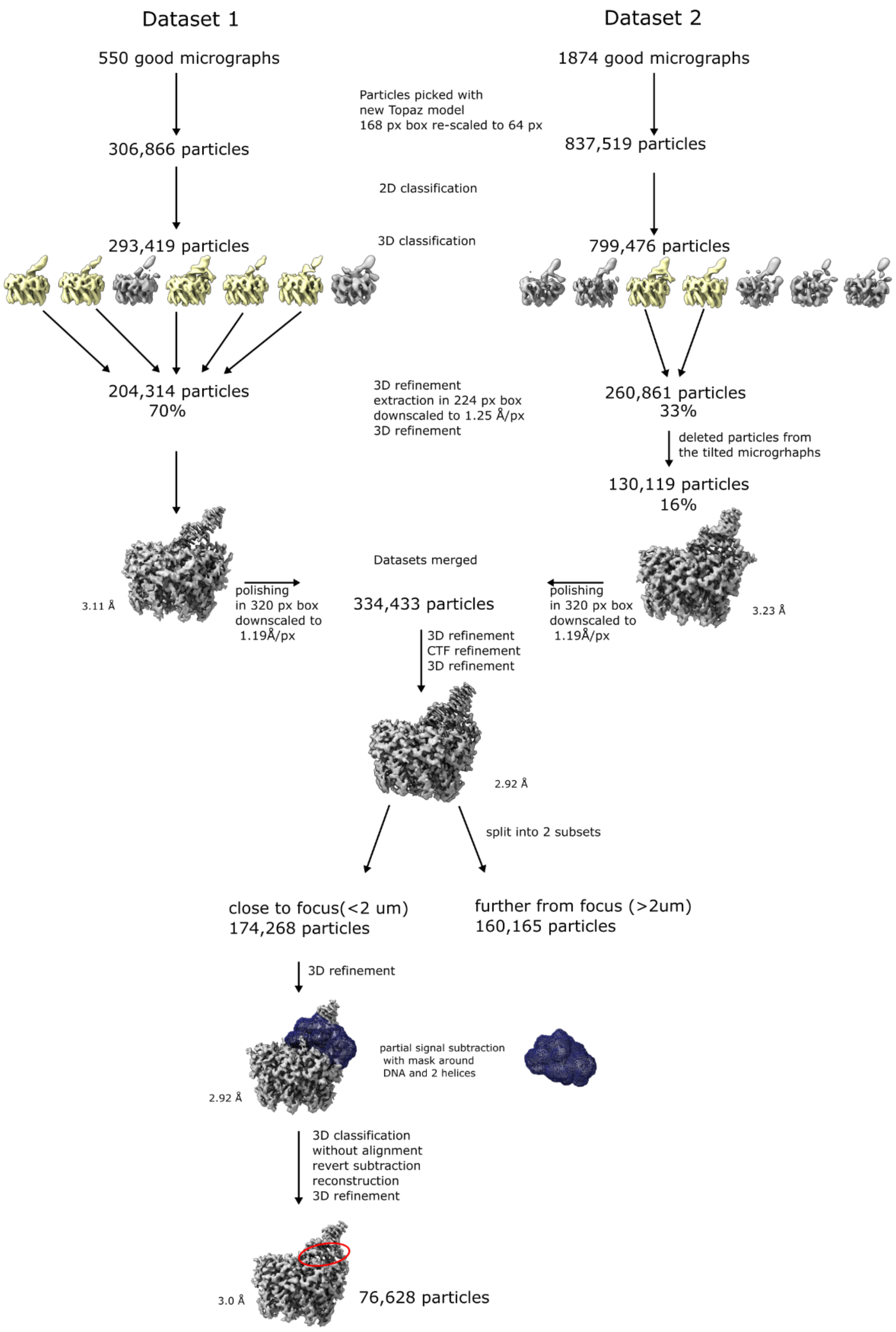

**Supplementary Figure 7**. **Final CryoEM processing flow chart for merged data from both datasets.** Overview of the cryoEM processing steps for the combined datasets. Particles were picked with a Topaz model trained with the "good particles" from the final model for dataset 2. Mask for focused classification with partial signal subtraction around two helices and a region of DNA is shown in blue. Red oval highlights the area of the structure where resolution was improved after focused 3D classification.

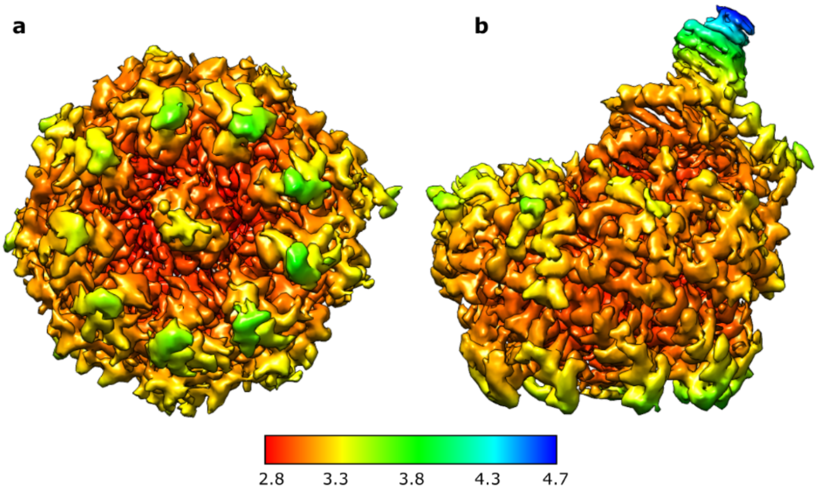

**Supplementary Figure 8.** CryoEM map coloured by local resolution. **a** Bottom view**. b** Side view.

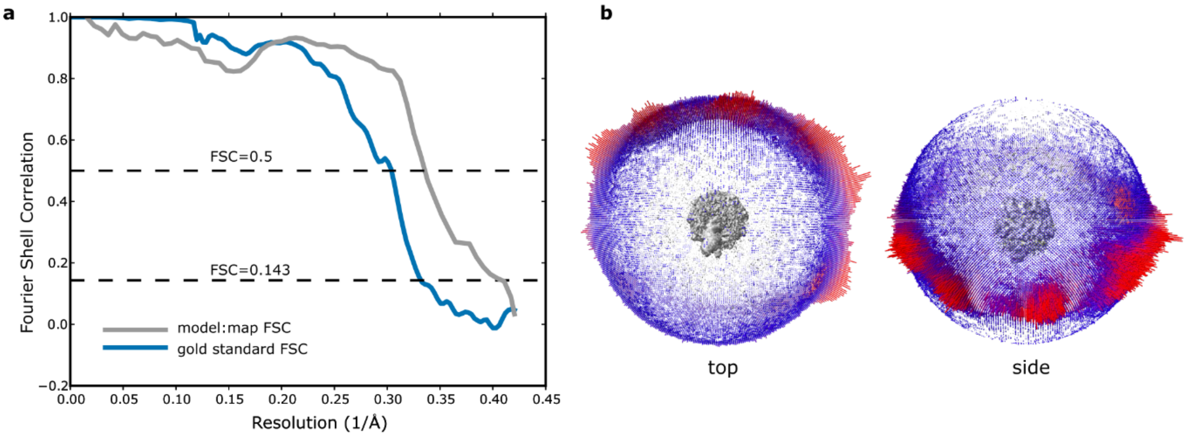

**Supplementary Figure 9. a** 'Gold standard' FSC curve (blue) and model:map FSC curve (grey). Dashed lines indicate FSC cut-offs of 0.143 and 0.5 for 'gold standard and model:map FSC, respectively. **b** Euler angle distribution of particles contributing to the cryoEM reconstruction.

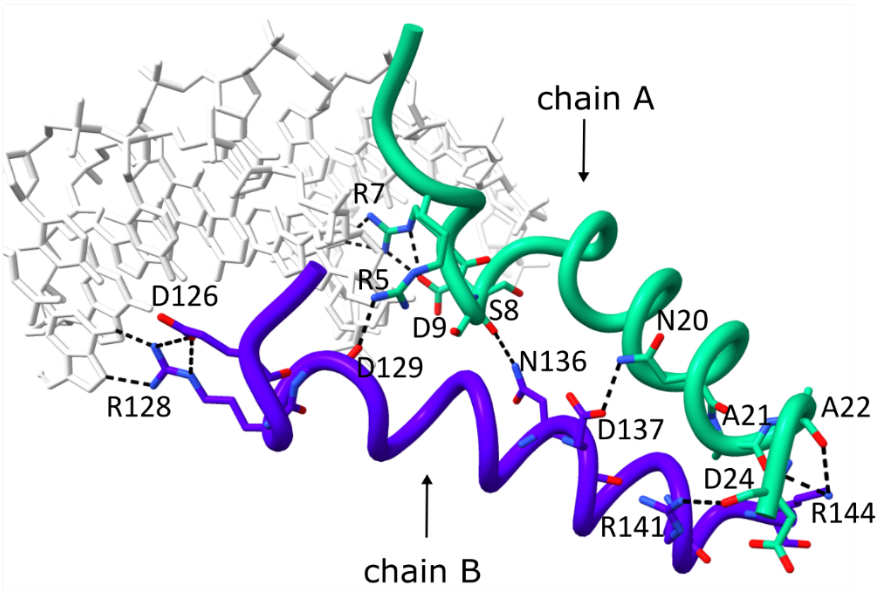

**Supplementary Figure 10. DNA binding substructure.** N-terminus of chain A is shown in green**,** C-terminus of chain B is in purple, DNA is in white, hydrogen bonds are depicted as black dashed lines.

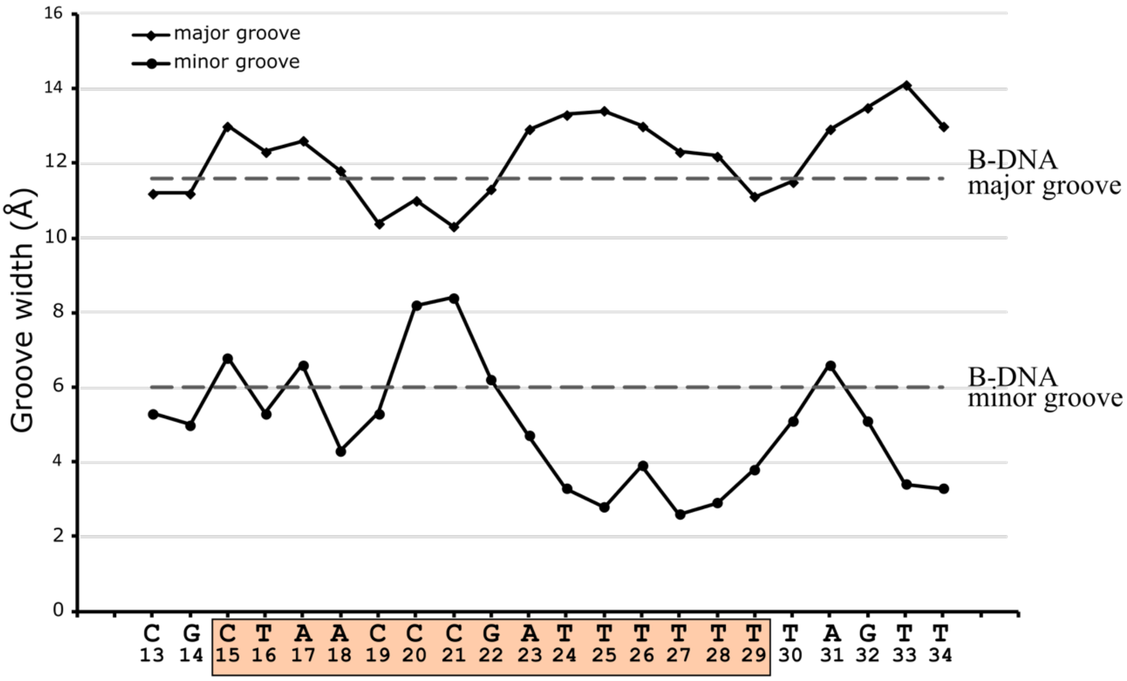

**Supplementary Figure 11. Major and minor groove width.** The widths of major and minor groove in DNA in the small terminase:DNA complex calculated using the program CURVES+ (Lavery & Sklenar, 1989). The average major and minor groove widths of B-form DNA (Chandrasekaran & Arnott 1996) are shown as a dotted line. Small terminase binding site is highlighted in orange.

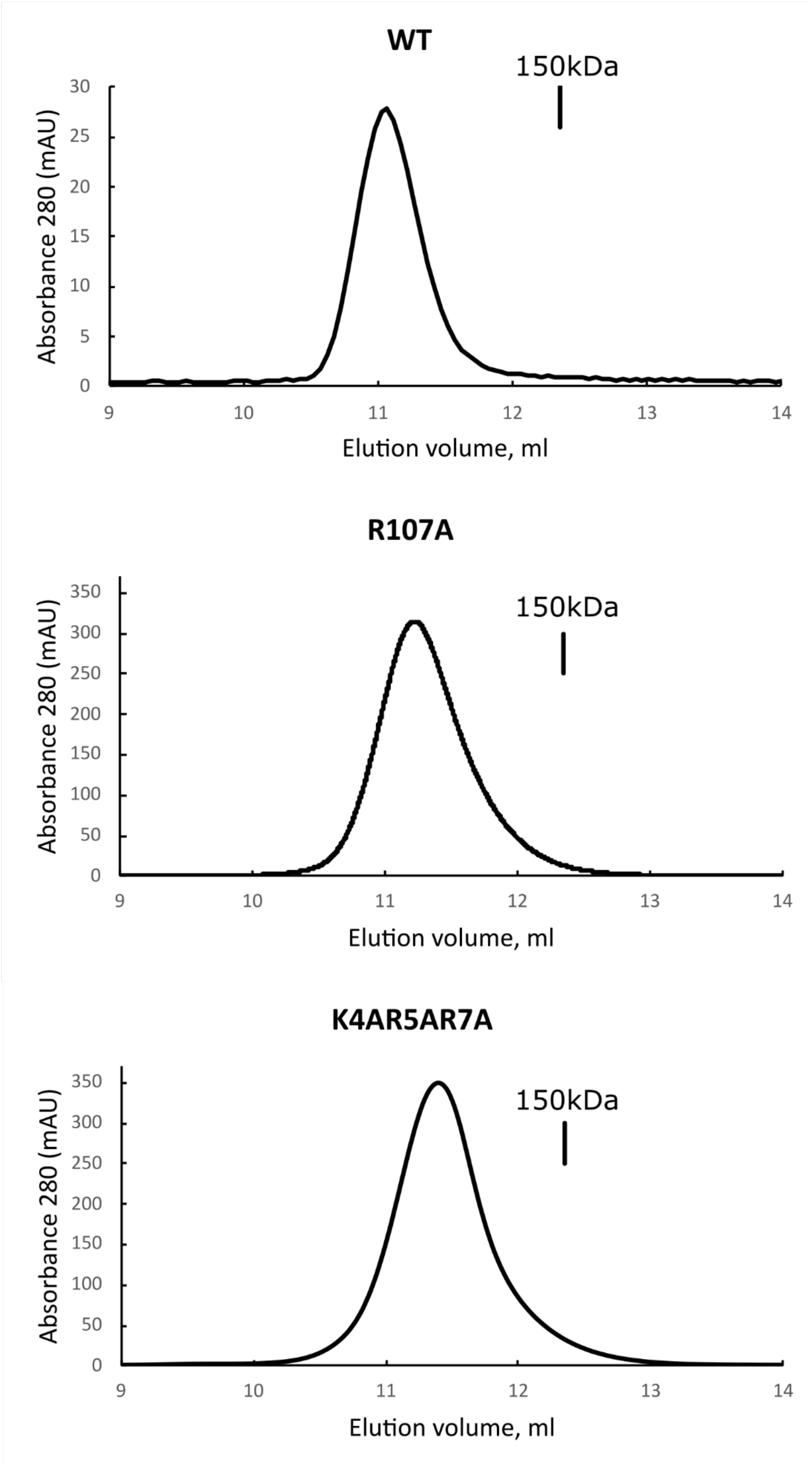

**Supplementary Figure 12. Purification of small terminase mutants.** Elution profiles of wild type (WT) protein and mutants R107A and K4AR5AR7A on S200 10/30 column.

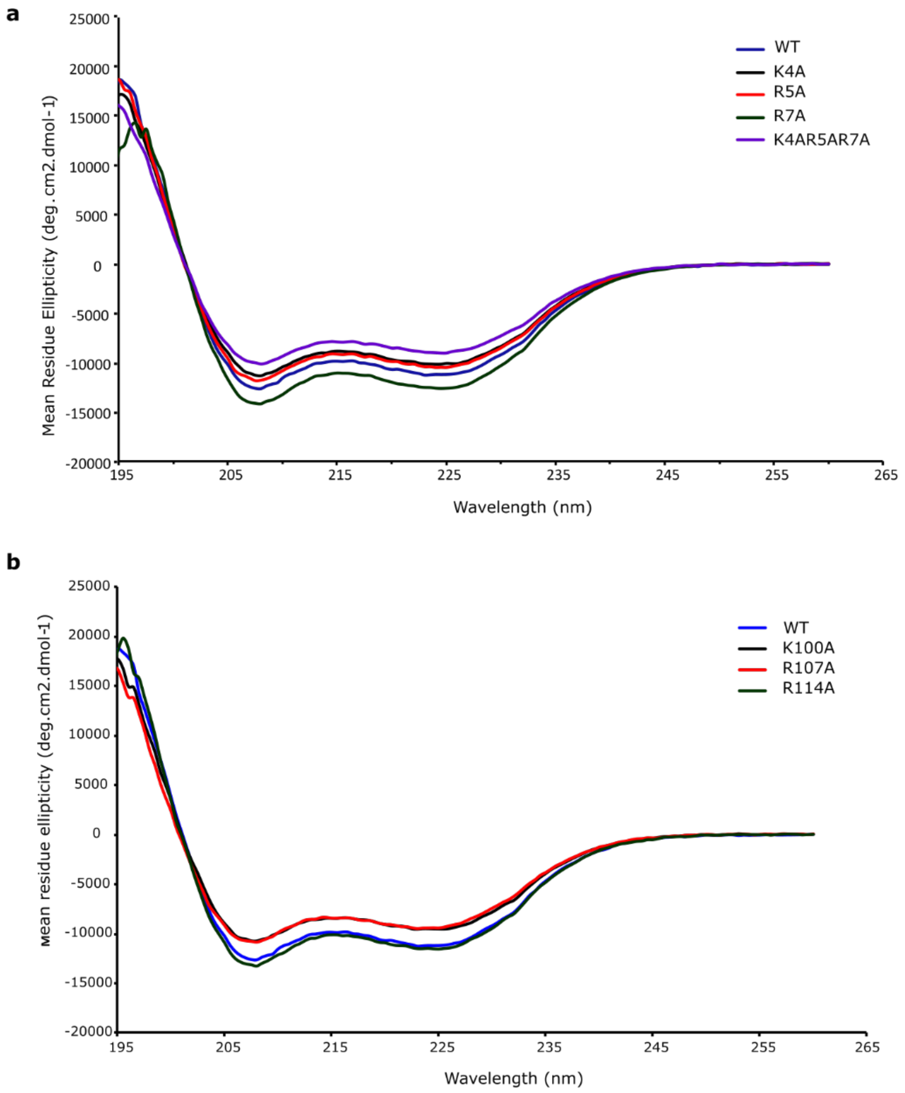

**Supplementary Figure 13. Far-UV circular dichroism spectra of wild-type (WT) and mutants of small terminase**. **a** Mutants of N-terminal arm. **b** Mutants within the channel of small terminase.

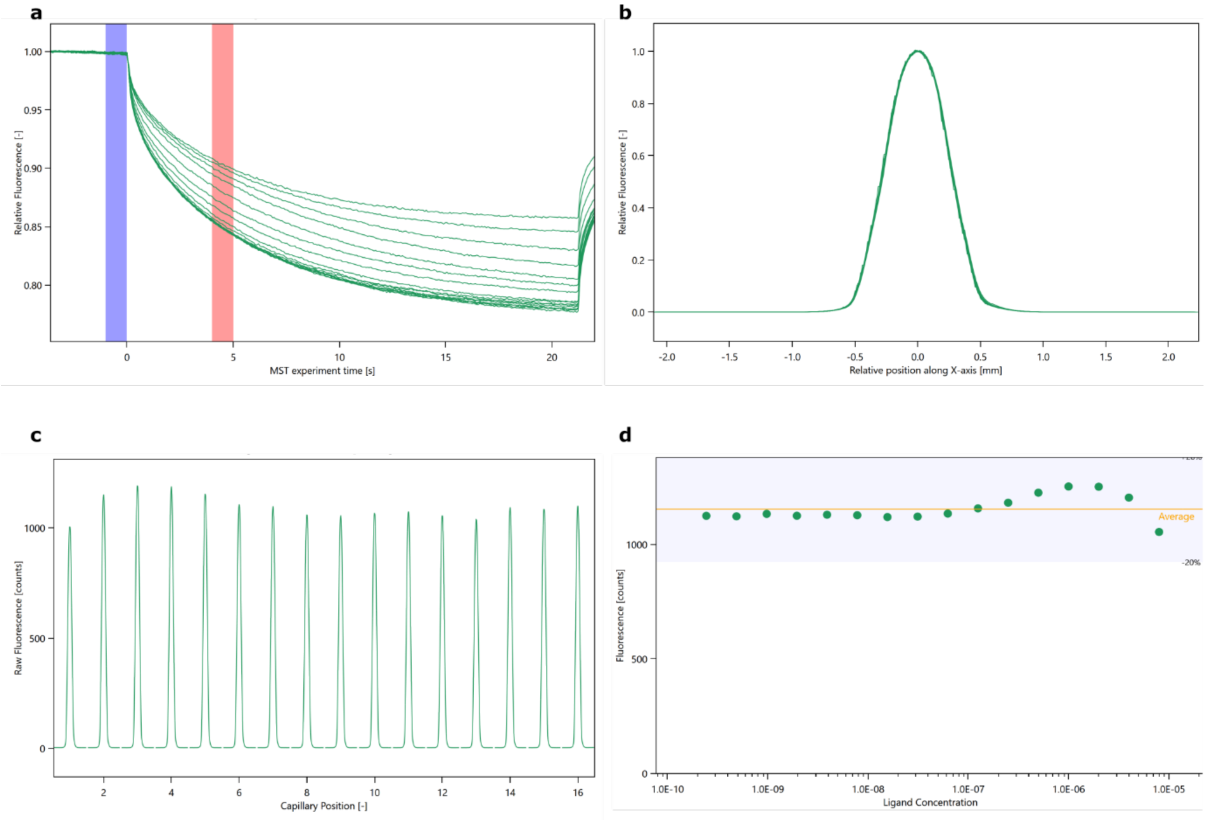

**Supplementary Figure 14. MST original data for one run of WT small terminase. a** Fluorescent traces over time for each capillary containing different concentrations of protein **b** Representative scan of fluorescence intensity across one capillary. **c** Representative capillary scan of fluorescent intensity across all capillaries. **d** Plot showing initial fluorescence intensity across all samples (capillaries) prior to temperature jump.

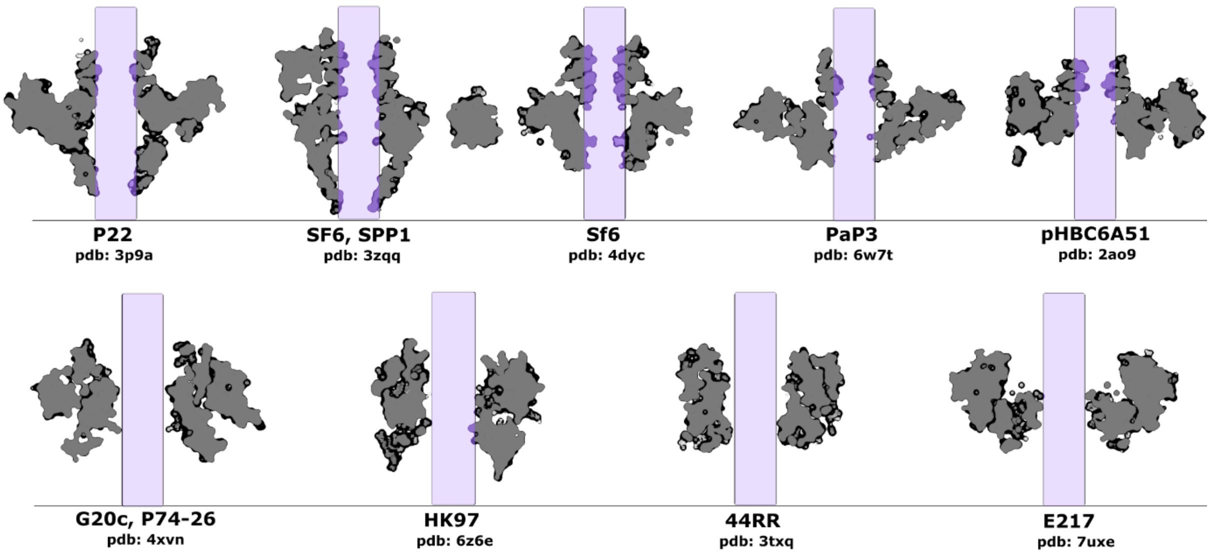

**Supplementary Figure 15. Comparison of the central channel of small terminases.** 3 Å thick section through the middle of small terminase models with 20 Å diameter cylinder positioned along the channel axis.

**Supplementary Table 1**. **Statistics for cryoEM data collection and processing**.

| Data collection | Dataset 1 | Dataset 2 |
|---|---|---|
| Microscope/detector | Krios/Gatan K2Summit with energy filter (slit width of 20 eV) | |
| Voltage (kV) | 300 | |
| Nominal magnification | 130,000x | |
| Recording mode | counting | |
| Flux (e- /Å$^2$/s) | 6.65 | 6.7 |
| Target defocus (µm) | 1.3 to 3.1 | 1 to 2.4 |
| Pixel size Å | 1.07 | 1.07 |
| Fluence (e- /Å$^2$) | 53 | 54 |
| Number of fractions | 50 | 50 |
| Total exposure time (s) | 8 | 8 |
| Number of movies | 682 | 2085 |
| Total particles picked | 306866 | 837519 |
| Particles from both datasets used in final reconstruction | 334433 | |
| Map resolution at FSC = 0.143 (Å) | 2.9 | |
| Local resolution (Å) | 2.9-4.9 | |
| Map sharpening B-factor (Å$^2$) | -49.9 | |
| Particles after focused classification with partial signal subtraction | 76626 | |
| Map resolution at FSC=0.143 (Å) | 3.0 | |
| Map sharpening B factor (Å$^2$) | -30.9 | |

**Supplementary Table2**. **Refinement statistics and validation**.

| | |
|---|---|
| Map resolution range refined against (Å) | 3.0 – 114.1 |
| Non-hydrogen atoms | |
| Protein | 7784 |
| Nucleic acid | 1145 |
| Average B factors (Å$^2$) | |
| Protein | 87.5 |
| Nucleic acid | 115.5 |
| R.m.s. deviations | |
| Bond lengths (Å) | 0.0059 |
| Angles (°) | 1.335 |
| Validation | |
| MolProbity score | 1.11 |
| Clashscore | 3.16 |
| Rotamer outliers (%) | 0.12 |
| Ramachandran plot | |
| Favoured (%) | 99.25 |
| Outliers (%) | 0.0 |

**Supplementary Table 3. N- termini sequences of DNA binding proteins with N-terminal arm (NTA) and all known small terminases.** Grey boxes- not modelled residues, light blue boxes – NTA, red font - positively charged residues, yellow boxes – non structured residues, green boxes – helices, purple box – β-sheet.

| PDB | Protein/phage | Sequence |
|---|---|---|
| 5ZJQ | Homeobox extradenticle chain A | KKRKPYSKFQTLELEKEF |
| 5ZJQ | Homeobox abdominal-B chain B | DARRKRRNFSKQASEILNEYFYS |
| 1W0T | hTRF1 | KRQAWLWEEDKNLRSGVRKYG |
| 1W0U | hTRF2 | KKQKWTVEESEWVKAG |
| HK97 | Enterobacteria phage HK97 | MADKRIRSDSSAAAVQAMKNAA |
| 4Z3C | Bacillus phage SF6 | MKEPKLSPKQERFIEEYFIN |
| 3HEF | Enterobacteria phage Sf6 | MATEPKAGRPSDYMPEVADDICSLLSS |
| 3P9A | Enterobacteria P22 | MAAPKGNRFWEARSSHGRNPKFESPEALWAAC |
| 2A09 | Bacillus cereus | MPFSISGRKGSEMMAKLDELKQKLTAK |
| 3TXQ | 44RR | MNDVLDFTQLKDLNGIEGIHGEDVQVYAPLVLRDPVSNPNNRKIDQDDDYELVRRN |
| 4XNV | Thermus phage G20c | MSVSFRDRVLKLYLLGF |
| 6W7T | Pseudomonas virus PaP3 | MSDEKVVSIGAAPLSAKEKLDLYCE |
| 7UXE | Pseudomonas E217 | MTKFYSPDDLVTPQEFADPHFAAINQKRFDLYIDLRVQG |
| 1J9I | Escherichia phage lambda | MEVNKKQLADIF |

**Supplementary Table 4.** Borders of DNA binding domains or N-terminus regions highlighted in gold in Supplementary Figure 1.

| Phage | PDB | |
|---|---|---|
| P22 | 3p9a | n-terminus: 4-24 aa |
| SP6 | 3zqq | HTH: 10-60 aa |
| Sf6 | 4dyc | HTH: 10-52 aa |
| PaP3 | 6w7t | HTH: 13-46 aa |
| pHBC6A51 | 2ao9 | HTH: 23-48 aa |
| G20c | 4xvn | HTH: 1-52 aa |
| HK97 | xxx | DBM: 3-24 aa chain A, 125-145 aa chain B |
| 44RR | 3txq | n-terminus: 25-40 aa |
| E217 | 7uxe | HTH: 14-50 aa |
